# Thymic epithelial cell fate and potency in early organogenesis assessed by single cell transcriptional and functional analysis

**DOI:** 10.1101/2023.06.07.544040

**Authors:** An Chengrui, Alison M. Farley, Sam Palmer, Dong Liu, Anastasia I. Kousa, Paul Rouse, Viktoria Major, Joanna Sweetman, Jan Morys, Andrea Corsinotti, Jennifer Nichols, Jan Ure, Renee McLay, Luke Boulter, S. Jon Chapman, Simon R. Tomlinson, C. Clare Blackburn

**Author notes:** Correspondence: C. Clare Blackburn. **Equal contribution and first authorship:** These authors contributed equally to this work and share first authorship (An Chengrui, Alison M. Farley, Sam Palmer). Walter and Eliza Hall Institute for Medical Research, Parvkville Victoria 3050, Australia (Alison M. Farley). Revvity, 8100 Cambridge Research Park, Waterbeach, Cambridge, CB25 9TL, UK. Memorial Sloan Kettering Cancer Center, New York, NY 10065, USA. Resolution Therapeutics, SCRM Building, 5 Little France Drive, Edinburgh EH16 4UU, UK. MRC Human Genetics Unit, Institute of Genetics and Cancer, University of Edinburgh, Edinburgh EH4 2XU.

## Abstract

During development, cortical (c) and medullary (m) thymic epithelial cells (TEC) arise from the third pharyngeal pouch endoderm. Current models suggest that within the thymic primordium most TEC exist in a bipotent/common thymic epithelial progenitor cell (TEPC) state able to generate both cTEC and mTEC, at least until embryonic day 12.5 (E12.5) in the mouse. This view, however, is challenged by recent transcriptomics and genetic evidence. We therefore set out to investigate the fate and potency of TEC in the early thymus. Here using single cell (sc) RNAseq we identify a candidate mTEC progenitor population at E12.5, consistent with recent reports. Via lineage-tracing we demonstrate this population as mTEC fate-restricted, validating our bioinformatics prediction. Using potency analyses we also establish that most E11.5 and E12.5 progenitor TEC are cTEC- fated. Finally we show that overnight culture causes most if not all E12.5 cTEC-fated TEPC to acquire functional bipotency, and provide a likely molecular mechanism for this changed differentiation potential. Collectively, our data overturn the widely held view that a common TEPC predominates in the E12.5 thymus, showing instead that sublineage-primed progenitors are present from the earliest stages of thymus organogenesis but that these early fetal TEPC exhibit cell-fate plasticity in response to extrinsic factors. Our data provide a significant advance in the understanding of fetal thymic epithelial development and thus have implications for thymus-related clinical research, in particular research focussed on generating TEC from pluripotent stem cells.

## Introduction

T cell repertoire development, called thymopoiesis, depends on dynamic interactions between the developing T cells (thymocytes) and the stromal compartment of the thymus (1, 2). This compartment is complex, comprising epithelial, mesenchymal, vascular and haematopoietic components organized into two regions, the cortex and the medulla, which support different aspects of thymopoiesis (3, 4, 5). Of the stromal elements, it is the thymic epithelium (TE) that directs most of the organ’s specialist functions (6). Thymic epithelial cells (TEC) comprise two functionally distinct subsets, cortical (c) and medullary (m) TEC. Broadly, cTEC regulate T cell lineage commitment and positive selection while mTEC impose central tolerance on the T cell repertoire (6, 7, 8, 9, 10, 11, 12).

During development TEC arise from the endoderm of the third pharyngeal pouches (3PP), which also generate the parathyroid glands. Strikingly, transplantation of 3PP alone is sufficient to direct thymus formation in an ectopic site, establishing that cTEC and mTEC both originate from progenitor cells in the 3PP (thymic epithelial progenitor cells; TEPC) (13, 14). When differentiation of the endodermal cells in the thymus domain of the 3PP is blocked, for instance by the lack of a functional allele encoding the transcription factor Forkhead Box N1 (FOXN1), a functional thymus able to support T cell lineage commitment and subsequent thymopoiesis cannot be generated (15, 16); FOXN1 is required throughout life for normal differentiation of both cTEC and mTEC (17) and is regarded as a master regulator of TEC differentiation (15, 16, 17, 18).

cTEC and mTEC are thought to arise from a common progenitor cell. Several studies have shown that the cTEC- and mTEC-sub-lineages are both generated from cells expressing the cTEC-associated markers CD205 and β5t (encoded by *Psmb11*) (19, 20). In addition, an elegant clonal analyses using single cell reversion of a null allele of *Foxn1* has shown that bipotent TEPC can exist *in vivo* (18), and a single cell transplantation analysis also suggested most cells in the early fetal thymus are bipotent TEPC (21). Based on these observations, a serial progression model of TEC differentiation has been proposed (22). This suggests that fetal TEPCs exhibit features associated with the cTEC lineage and that additional cues are required for mTEC specification from this common TEPC (22).

Testing this model has been hampered by difficulties related to identification of cTEC-restricted sub-lineage specific progenitor TEC in the fetal thymus, due to shared expression of surface antigens between this presumptive cell type and the presumptive common TEPC (20, 22, 23), although cTEC-restricted progenitors clearly exist in the postnatal thymus (24). In contrast, the presence of functionally validated mTEC-restricted progenitors has been detected from day 13.5 of embryonic development (E13.5) (25). Fetal mTEC progenitors are characterized by expression of Claudin3/4 (CLDN3/4) and SSEA1 (26, 27). A CLDN3/4^high^ subpopulation was observed in the E11.5 thymic primordium (26) and CLDN3,4^high^SSEA-1^+^ TEC were present in *Foxn1^-/-^* thymi by E13.5 (17, 28), suggesting that divergence of cTEC and mTEC fates can occur independently of FOXN1. However, lineage divergence can also occur downstream of FOXN1 expression (18).

Receptors leading to activation of the Nuclear Factor kappa-light-chain-enhancer of activated B cells (NF-κB) pathway, including Lymphotoxin-β receptor (LTβR) and Receptor Activator of NF-κB (RANK), regulate the proliferation and maturation of mTEC (29, 30, 31) and a hierarchy of intermediate progenitors specific for the mTEC sub-lineage has been proposed based on genetic analysis of NF-κB pathway components (28, 32). In this model, non-canonical NF-κB signalling through RELB leads to an upregulation of RANK in early pro-mTEC cells but does not affect the generation of SSEA1^+^ progenitor mTEC, which are present in *Relb^-/-^*mice (28). Related, it has recently been shown that Notch signalling is essential for development of the earliest CLDN3/4^+^ mTEC progenitors in the fetal thymus, and that all mature mTEC, but not cTEC, have received high Notch stimulation (33, 34). In these experiments, abrogation of Notch signalling at ∼E11.25-E12 (using a *Foxn1Cre;Rbpj^fl/fl^* model) resulted in a less severe medullary phenotype than blockade of Notch signalling in the endoderm from E9.0 using *Foxa2Cre;dnMAML^fl/fl^* mice; in the latter model, CLDN3^+^ mTEPC were completely absent from the developing thymus at E14.5 and no mTEC development could be detected, at least until E16.5 (33). These data established that the earliest mTEPC must arise prior to the onset of *Foxn1Cre* activity, namely, prior to ∼E11.25. Therefore, consistent with phenotypic analyses (17, 26, 28), but in contradiction to the serial progression model, they predict the presence of mTEC-fated TEC within the thymic primordium prior to E12.5.

Early thymus organogenesis has recently begun to be mapped using scRNA sequencing (35, 36) and *in vivo* barcoding (37). In particular, Magaletta and colleagues found that TEC could be clustered into two populations at E12.5: a large cluster the authors called “cTEC” and a smaller *Plet1^+^Cldn3,4^+^* cluster, “mTEC” (36). Furthermore, *in vivo* barcoding initiated by *Foxn1Cre* supported the presence of cTEC-fated TEPC in the fetal thymus but did not reveal a common or mTEC-restricted progenitor active in the fetal thymus; indeed, this study did not provide information regarding the source of mTEC during fetal development, although both mTEC-biased and common TEC progenitor activities were observed postnatally (37). Collectively therefore, these data also support a model in which cTEC and mTEC fates initially diverge early in thymus organogenesis.

To provide clearer understanding of the early events in TEC lineage progression, we set out to examine the fate and potency of TEC in the early thymic primordium using a combination of scRNAseq, lineage tracing and functional analyses. Here, using scRNAseq, we confirm the presence of candidate mTEC progenitors at E12.5 and identify potential transcriptional regulators of this and other E12.5 TEC populations. We then use a combination of lineage-tracing and potency analyses to address the fate and potency of E12.5 TEC populations, establishing that at E12.5 the majority of progenitor TEC are cTEC-lineage restricted while a minor population of mTEC-restricted TEC exists by this time point. Finally, we show that overnight culture changes the potency of E12.5 cTEC-restricted TEPC such that most/all of these cells become bipotent, and reveal possible mechanisms for this change in potency through scRNAseq. Collectively, our data overturn the widely held view that a common TEC progenitor predominates in the E12.5 thymus, showing instead that sub-lineage-primed progenitors arise from the earliest stages of thymus organogenesis. Reconciling these two models, they also establish the plasticity of early fetal TEPC in response to extrinsic factors. Our data provide a significant advance in understanding fetal thymic epithelial development and have implications for understanding lineage relationships in later development and the postnatal thymus.

## Materials and Methods

### Mice

C57BL/6 mice were used for isolation of fetal TEC except for the single cell transfer experiments, which used C57BL/6 x CBA F1 embryos. For timed matings, noon of the day of the vaginal plug was taken as day 0.5 of embryonic development (E0.5). Foxn1^GFP^ (38), Foxn1^Cre^ (39), Sox9CreER^T2^ (40), Gt(ROSA)26Sor^tm14(CAG-tdTomato)Hze^ (Ai14) (41), Rosa26*^CAG-STOP-Foxn1-IRES-GFP/+^* (iFoxn1; (42)) and Rosa26-CreERT2 (43) mice were as described. For the clonal assay experiments we used a Cre-inducible membrane-bound EGFP reporter mouse strain (mGFP) that directs GFP expression to the cell membrane and which expression is present in the majority of cells of all tissues. This reporter mouse model and design is as described by Gilchrist and colleagues (44), except that the EGFP has been modified with the addition of a RAS farnesylation sequence, and was a kind gift from Alexander Medvinsky (University of Edinburgh). Adult male C56BL/6xCBA F1 mice were used as graft recipients except where otherwise noted. All animals were housed and bred in University of Edinburgh animal facilities. All experimental procedures were conducted in compliance with the Home Office Animals (Scientific Procedures) Act 1986, under project licences PPL60/3715 and PEEC9E359 to V. Wilson. Primers used for genotyping were as shown in Supplementary Table 1. All controls were littermates unless otherwise stated.

### Tamoxifen administration

Tamoxifen was administered to pregnant female mice at the ages noted. Briefly, 300mg tamoxifen (Merck, PHR2706) was suspended in 1.2ml EtOH, vortexed for 5 mins, placed at 37°C for 30 minutes, then suspended in pre-warmed (42°C) corn oil to a final concentration of 40mg/ml. Female mice at E13.5 were then treated by oral gavage with a single dose of 16mg tamoxifen per mouse. Female mice at E11.5 were treated by gavage with two doses of 8mg tamoxifen at 24-hour intervals. The female mice were sacrificed for embryo collection on either E13.5 and E18.5 (after tamoxifen administration at E11.5 and E12.5), or E18.5 (after tamoxifen administration at E13.5).

### Thymus dissociation

For single cell transfer experiments, microdissected fetal thymic lobes were resuspended in 0.7mg/ml hyaluronidase, 0.35mg/ml collagenase, 0.05mg/ml deoxyribonuclease in PBS for 15 minutes at 37°C. Gentle pipetting was used to aid dissociation and the cells were then spun down and collected in FACS wash (PBS with 5% fetal calf serum [FCS]). For all other experiments, microdissected fetal thymi were dissociated for 5 minutes in TrypLE Express Enzyme (Life Technologies 12604013) in an Eppendorf Thermomixer (1400 rpm, 37°C) and were then triturated with a P1000 and the 25G syringe ten times each. Cell suspensions at 4°C were washed twice in 2% FCS FACS buffer, resuspended as required and filtered through a 70μm cell strainer (Corning) to remove clumps.

### Flow Cytometry

Cells were processed for flow cytometric sorting and analysis as previously described (17, 42). Compensation controls were performed using beads. Sorting and analysis gates were set using FMOs. Sorting was performed using a MoFlo MLS high-speed sorting flow cytometer (Beckman Coulter), BD FACS Aria II or Fusion running FACS Diva 4.1 (BD Biosciences). FACS analysis were performed on a FACSCalibur (BD Bioscience) or Novocyte running NovoExpress 1.3.0 (ACEA) at the CRM, University of Edinburgh. All post-acquisition analysis was performed with FloJo (Tree Star Inc) or FCSexpress 6 (De Novo Software) software.

### Immunohistochemistry

For Figures 4-6, immunohistochemistry was performed as described (14). For analysis of E18.5 samples, thymi were taken into sucrose before sectioning. Appropriate isotype and negative controls were included in all experiments. Images were captured using a Leica AOBS or SP8 confocal microscope and processed using Adobe Photoshop CS2 (Figures 4-6) or Leica LASX (Figures 2, 3) software. Images presented are of single optical sections. Videos were made in MATLAB. For Figures 2, 3, immunohistochemistry was performed on thick sections as follows: after sacrificing the embryos, they were fixed overnight in 4% paraformaldehyde, then washed three times with PBS for one hour. Samples were then embedded in 1% agarose and 200μm sections were cut using a vibratome. Sections were cleared using X-Clarity^TM^ (Logos Biosystems). Briefly, the X-Clarity protocol was as follows: the vibratome sections or sample were washed with PBS then incubated with Hydrogel Solution (Logos Biosystems. C13103) for 24 hours at 4℃. The samples were then polymerized in Hydrogel Polymerization System (Logos Biosystems) for 3 hours, washed three times in PBS for one hour each, then incubated in Electrophoretic Tissue Clearing Solution (Logos Biosystems C13001) overnight. They were then washed in 1% Triton X100 in PBS (PBST) for one hour, three times. Sections were then stained as follows: sections were blocked in 1% goat serum then incubated with the appropriate primary antibody solution at room temperature for two days, then washed in 1% PBST for one hour, three times. Sections were then incubated in 1:1000 secondary antibody and DAPI (1:1000) solution at room temperature for one day and washed in 1% PBST for one hour. Steps after blocking was repeated if further antibody staining was required. The same staining protocol was followed for whole-mounted samples (Figure 3).

### Antibodies

The antibodies used for immunohistochemistry and flow cytometry were as listed in Supplementary Table 2.

#### Single cell RNA-seq

##### Smart-seq 2

Individual EPCAM^+^PLET1^+^ cells from E10.5 or E12.5 primordia were sorted into 96-well BD Precise WTA Single Cell Encoding Plates (see Supplementary Figure 1 for sorting strategy). The plates were sealed and stored at -80°C until library preparation. Four plates (376 cells) were collected per timepoint. Reverse transcription was performed in the presence of ERCC RNA control. The 96 samples from each plate were then pooled and purified with Agencourt® AMPure® XP magnetic beads (clean-up step). Second strand synthesis was carried out and the resulting products cleaned up. Following adaptor ligation and clean-up, ligation products were PCR amplified for 18 cycles overnight, and all subsequent steps were performed in a separate post-amplification area. The amplified products were cleaned up and quantified using Qubit dsDNA HS Assay. 50ng of the products then underwent random primer extension and 12 cycles of further amplification. The amplified libraries were cleaned up and quantified with Qubit assay. The libraries were stored at -80°C until sequencing. The libraries were sequenced on Illumina MiSeq at the Wellcome Centre for Human Genetics in Oxford within 6 months of preparation.

Pre-processing of the FASTQ files was performed using the BD: Precise Whole Transcriptome Assay Analysis Pipeline v2.0. This pipeline performs all the required steps to demultiplex, align, and quantitate sequencing reads from BD™ Precise Whole Transcriptome Assay. After performing a quality control assessment using FastQC (45), the pipeline filters and demultiplexes sequencing reads based on the sequence of the 8 base sample barcode (46). It aligns reads from each sample to the genome using STAR (47), with feature counting by the union rules of HTSeq-count (48). The initial quantification is performed by counting the unique molecular indices (MI) mapped to each feature (49). Lastly, the pipeline uses two error correction algorithms to remove MI errors. In brief, MI errors that are derived from sequencing base calls and PCR substitution errors are identified and adjusted to a single MI barcode using Recursive Substitution Error Correction™ (RSEC). Subsequently, MI errors that are derived from library preparation steps or sequencing base deletion errors are adjusted using Distribution-Based Error Correction™ (DBEC). Following this pre-processing, transcripts from the same gene were combined using the maximum values; protein coding genes were selected; cells with more than 50% mitochondrial genes were deleted; cells with reads number between 40000 to 400000 and features between 500 to 5000 were selected; and CPM was calculated. 6: genes with CPM>2 at least in 2 cells were selected. The data were then analysed using Seurat (v3.1.4) in R (v3.6.1). To reduce the batch effect between plates in library construction, t-SNE transformation was used to normalize the data. The remaining steps, including feature selection, principal analysis, and K-Nearest Neighbour clustering, were as set out in Seurat. Regulon analysis was performed with SCENIC (v1.2.1.1) according to its guidelines (50).

#### 10X protocol and analysis

##### Experimental design

E12.5 and E13.5 wild type (WT), E12.5 and E13.5 Rosa26CreERt2;iFoxn1 (iFoxn1; called enforced Foxn1) and E12.5 and E13.5 Rosa26CreERt2 (called Cre-only) fetal thymi were microdissected and genotyped prior to further processing. The samples processed for library preparation were as follows: (i) Freshly isolated wild type E12.5 and E13.5 and E12.5 iFoxn1 and Cre-only lobes were dissociated to single cell suspensions using TrypLE Express (Life Technologies 12604013) as above and immediately processed for library preparation; (ii) E12.5 wild type, iFoxn1 and Cre-only lobes were cultured overnight as intact lobes at the air-liquid interface (i.e. standard fetal thymic organ culture conditions, called FTOC), then processed to single cell suspensions using TrypLE Express; and (iii) E12.5 wild type, iFoxn1 and Cre-only lobes were dissociated to single cell suspensions and cultured overnight as monolayers (called monolayer), then harvested using TrypLE Express. Culture was in Advanced DMEM/F-12 (Gibco 12634010), 2% FCS, 1% Pen/Strep, 1% GlutaMAX, 1% non-essential amino acids.

##### Processing of samples for library preparation, library preparation and sequencing

Cells from each condition were resuspended in 90μl FACS buffer (PBS without Ca^2+^ or Mg^2+^ with 10% FCS), stained with barcode tagged antibodies against MHCII (B0117), CD40 (B0903), CD80 (B0849), with biotinylated UEA1 bound to SAV-PE-B0952) and with α-EPCAM-magnetic beads (Miltenyi) (see Supplementary Table 3), and washed. Each condition was then hashed with 100ul lipid anchored Cell Multiplexing Oligos for sample multiplexing (CMOs, 10x Genomics). The CMOs for each sample were as follows: CMO 301, enforced Foxn1 FTOC; CMO 302, Cre-only FTOC; CMO 303, WT FTOC; CM304, enforced Foxn1 monolayer; CMO5, Cre-only monolayer; CMO6, WT monolayer; CMO7, enforced Foxn1 E12.5; CMO8, Cre-only E12.5; CMO9, WT E12.5; CMO10, enforced Foxn1 E13.5; CMO11, Cre-only E13.5; CMO12, WT E13.5). Samples were then combined and passed through a LS Column (Miltenyi) to enrich for EPCAM^+^ cells. Cells eluted from the column were spun down and resuspended in 50µl FACS buffer, of which 10µl was taken for cell counting. Approximately a total of 15000-30000 cells pooled from each condition were then filtered and loaded on a Chip G (10X Genomics). Cell-bead encapsulation was performed running the chip in a Chromium X instrument (10X Genomics). Upon encapsulation, gene expression and multiplexing libraries were prepared following the manufacturer’s instructions using the Chromium Next GEM single cell reagent kit 3’ V3.1 (dual index) with Feature Barcoding technology for Cell Multiplexing (CG000388, 10x Genomics). Libraries were quantified using a Bioanalyzer DNA High-sensitivity Kit (Agilent Technologies), pooled at a molar ratio of 6:1 between gene expression and multiplexing libraries, and sequenced on an Illumina NextSeq 2000 P3 flow cell (100 cycles configuration) to obtain approximately 1.4B paired-end reads.

##### QC and analysis

FASTQ files were processed using Cell Ranger (10x Genomics) and then count matrices (for RNAseq, CITEseq and CMO multiplexing) were analyzed using the R package Seurat. Data from three experimental time points (E12.5, E13.5 and overnight culture) were processed separately and then combined. First, cells were filtered to include only cells with more than 200 RNA features. Then Seurat’s *SCTransform* function was applied, regressing on percentages of mitochondrial transcripts. Then clustering was performed using 30 PCA dimensions with resolution 0.15. For each time point, there were two clusters corresponding to TEC, with the other clusters corresponding to thymocytes (marker *Tcrg-C1*), fibroblasts (marker *Prrx2*), neural cells (marker *Neurod1*), parathyroid cells (marker *Gcm2*) or cells with ribosomal RNA contamination (marker *Gm42418*). There was also a small cluster excluded at this stage with high counts for the multiplexing CMO IDs CMO304-CMO306, corresponding to the monolayer experimental conditions. The remaining six TEC clusters were then combined into one Seurat object and *SCTransform* was applied again, regressing on percentages of mitochondrial transcripts. Once again, small clusters with markers *Prrx2* and *Neurod1* were excluded. Finally, UMAP dimensional reduction and clustering was performed using 50 PCA dimensions and resolution 0.07 (after setting an RNG seed of 100).

Cells from the three time points (E12.5, E13.5 and overnight culture) were easily distinguished because they came from separate 10x runs, but within each time point, multiplexing was used to separate different experimental conditions. For the multiplexing analysis, CMO tags were used to distinguish between cells with enforced Foxn1 (iFoxn1), Cre only or wild type (WT) and also between cells from FTOC or Monolayer for the cells from the overnight culture condition as described in Materials and Methods (see also Supplementary Figure 2). Unfortunately, some of the CMO tags were very low (Supplementary Figure 2) and therefore all cells from the overnight culture condition (with CMO tags CMO301-CMO306) were originally assigned to identity CMO301 by Cell Ranger. We therefore overrode the CMO identity in cells with more than 20 counts for either CMO302 or CMO303, or with more than 2 counts for either CMO304, CMO305 or CMO306, with the corresponding CMO ID. CMO identity appeared to have only a modest effect on overall gene expression (Supplementary Figure 2), perhaps due to FOXN1 saturation or enforced FOXN1 supressing endogenous *Foxn1* expression (Supplementary Figure 2). We therefore ignored CMO identity in the analysis in the main text.

GO enrichment analysis was performed in R with the enrichR package, using the “GO_Biological_Process_2021” database. The Seurat function *DEenrichRPlot* was used with “max.genes=1000”. DE gene analysis was carried out in R using the Seurat function *FindMarkers*.

### Cell potency assays

Fetal TEC of the desired age were obtained by microdissection, dissociated to single cell suspensions as above and placed on a tissue culture treated 10cm Petri dish in a drop of media so that single cells could be observed for selection. In a neighbouring drop, E12.5 wild type lobes were placed ready for injection. Paraffin oil was poured around the drops to keep them in place. Using a microinjecting microscope and rig, pulled glass pipettes were manipulated by hand controllers to select the desired number of viable cells and inject these into the wild type lobes. All cells and injected lobes were kept on ice. The injected lobes were either transferred into media before grafting on the same day or were placed in Terasaki plates (Nalge Nunc International) in a drop of media and cultured overnight before grafting the next morning. Where noted, cells or lobes were cultured over night at 37°C at 7% CO_2_ in DMEM, 10% FCS, before grafting the next day.

### Kidney Capsule Grafting

The surgical operations were carried out on C57BL/6xCBA Fl male mice in a laminar flow hood under sterile conditions using a dissecting microscope (Olympus SZ40) as previously described (51). A small tear was made in the kidney capsule using fine forceps and the lobe was placed under it using a pulled glass pipette controlled by mouth suction.

### Tissue Culture

All cell manipulations were performed in a laminar flow sterile hood using sterile technique. Cell culture plastic ware was supplied by Iwaki. All solutions were tested for sterility and warmed to 37°C prior to use. Cells were examined using an inverted microscope (Olympus CK2).

### Statistics and experimental design

For all scRNAseq analyses, n represents the number of independent biological experiments. For cell potency assays, n represents an independent graft; at least three biologically independent replicates were performed for each injection condition. No statistical method was used to predetermine sample size, the experiments were not randomized, and the investigators were not blinded to allocation during experiments and outcome assessment. There were no limitations to repeatability of the experiments. No samples were excluded from the analysis, however, for the cell potency analyses only grafts in which mGFP^+^ cells were observed upon visual inspection under a fluorescence dissection microscope were analysed by immunohistochemistry.

### Statistical models

To estimate the proportion of cells that are cTEC fate-restricted, *p*, we analysed the transplantation assay results using two models representing extreme cases of maximally segregated spatial patterning (Model 1) and a well-mixed scenario (Model 2). In model 1, we assume that the number of cells taken, *N,* is small relative to the size of the thymus and the size of the cTEC regions. In this model, when cells are taken, they either come from a cTEC region and are all cTEC fated, with probability *p*, or they come from a region with mixed cell types and give rise to both cTEC and mTEC, with probability 1-*p*. In this model we don’t need to estimate the survival probabilities of cells, we can just take the number of experiments where some cells survived (n=23) and count the number of trials with only cortex contributions (x=17), see Table 4. This gives a binomial distribution for *p* with x=17 and n=23, resulting in an estimate of p=17/23=74% with a 95% C.I. of 56-92%.

In model 2, we assume each cell has an *independent* probability *p* of being cTEC-fated and we let *N* be the number of cells that survive the transplantation process. When small numbers of cells are transplanted, the survival rate is very small (approximately 1/83, see Table 4), which would suggest that in experiments where multiple cells are taken, only approximately *N=1* cell would survive, which leads to the same estimate and C.I. for *p* as model 1. If more than *N*=1 cell survive, then the estimate and C.I. for *p* become closer to one. For example, suppose 60 cells are taken and *N=*4 cells survive, let X be the random variable representing all cells being cTEC fated, such that

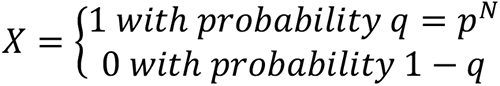

Since five out of eight grafts had only cortex contributions, the expectation of q is 5/8 with a 95% C.I. of 0.29-0.96, corresponding to an expectation that *p*=*q*^1/N^=89% with a 95% C.I. of 73-99%. If the number of surviving cells is even higher at *N*=10, we get *p*=95.4% with a 95% C.I. of 88-99.6%. In the experiments with overnight culture, when 30-40 cells were injected into recipient lobes, cultured overnight and then transplanted, one out of four grafts had only cortex contributions. When 1 cell was injected, zero out of four surviving grafts had only cortex contributions. This strongly suggests that model 2 (with a high number of surviving cells) is not suitable, and that either model 1 is appropriate, or equivalently model 2 with only *N*=1 surviving cell. In this case, since there were eight grafts in total with one having only cortex contributions, we estimate the proportion of cTEC-fated cells as *p*=1/8 with a 95% C.I. of 0-35%. We can compare the proportion of cTEC-fated cells in the original experiment (i.e. mGFP^+^ TEC-injected thymic lobes grafted with no overnight culture) with the overnight culture experiment (i.e. mGFP^+^ TEC-injected thymic lobes grafted after overnight culture) using a pooled two-proportion z-test to get a z-score of z=3.03 and a p-value of p-val=0.002.

### Data availability

scRNAseq data are deposited in GEO and are available through the following links: E10.5 and E12.5 TEC: GSE232765, Effects of overnight culture on TEC: GSE229248. The codes for the analyses shown in Figures 1-3 and 7 and related supplementary data are available at: https://github.com/purewaltzan/ThymusDevelopment.

## Results

### Single cell transcriptome analysis identifies discrete subpopulations among thymic epithelial cells in early organogenesis

Recently, we showed that Notch signalling is required for establishment of the mTEC lineage, and that this requirement is first evident prior to E12.5; in *FoxA2^Cre^;dnMAML^fl/fl^* mice, in which deletion occurs in the endoderm from ∼E9.0, CLDN3/4^+^ cells were absent from the early thymus primordium (33) and in *Foxn1^Cre^;Rbpj^fl/fl^* mice, in which deletion occurs in TEC from ∼E11.25, the mTEC compartment was significantly smaller than controls (33, 34). This suggested that the mTEC sublineage was specified prior to the onset of *Foxn1^Cre^*activity, and therefore that the mTEC-fated cells should be identifiable among TEC at E12.5 and possibly earlier stages, through single cell analyses.

To test this idea, we established scRNAseq libraries from TEC isolated from the E10.5 3PP and E12.5 thymic primordium, using the BD Precise WTA Single Cell Encoding Plate methodology (Figure 1A). The resulting dataset comprised 709 single-cell transcriptomes (352 from E10.5 and 357 from E12.5), with a median of 2318 unique genes per cell. Unsupervised clustering using Seurat revealed eight clusters within the dataset (Figure 1). Clusters 1-3 predominantly contained E10.5 cells, Cluster 4 contained both E10.5 and E12.5 cells and Clusters 5-8 contained predominantly cells from E12.5 (Figure 1B, C). The top ten differentially expressed marker genes (DEG) for each cluster were calculated using Seurat and visualized with a heatmap, with selected markers also visualized using a violin plot (Figure 1 D, E).

**Figure 1:**
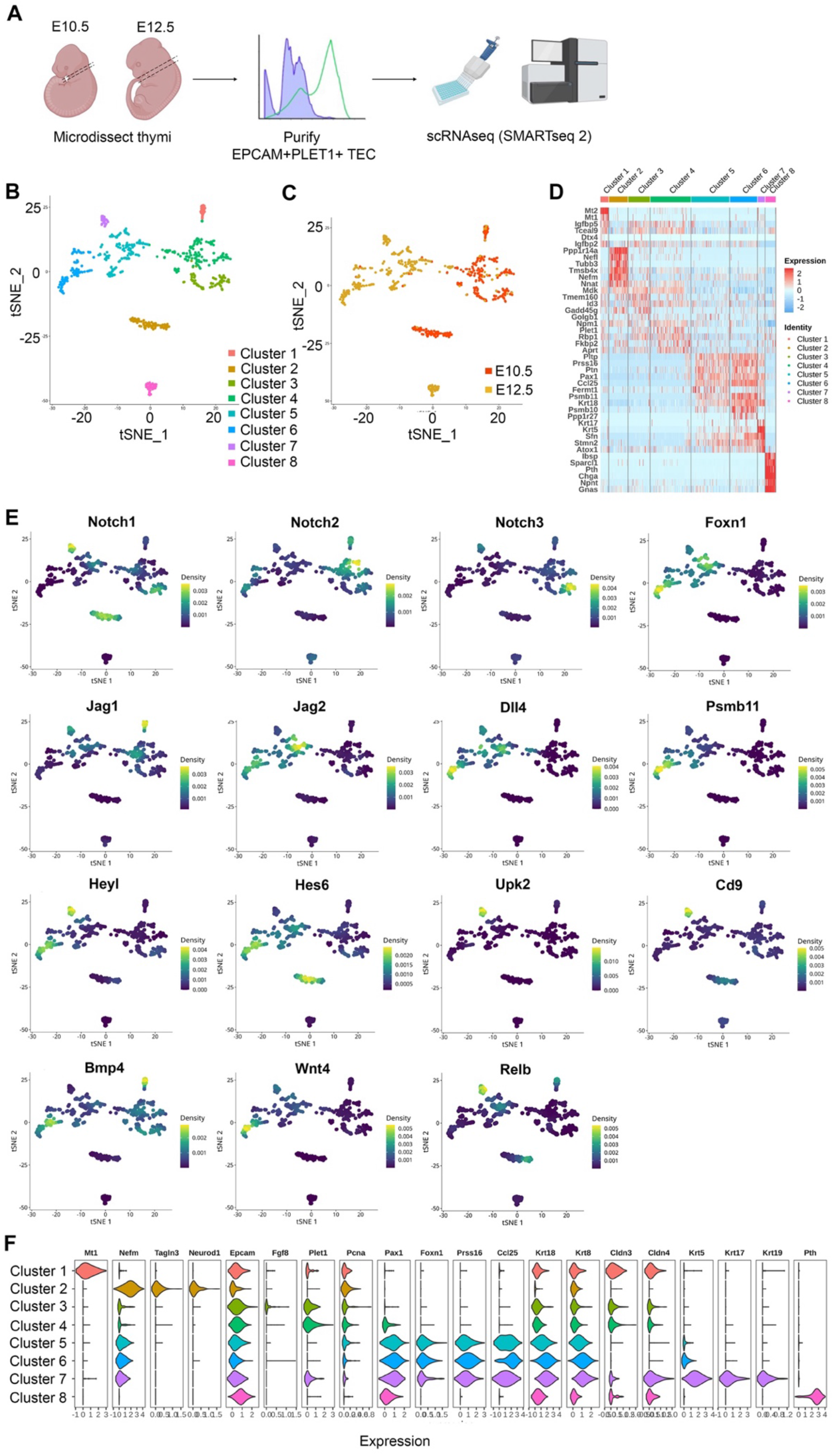
Heterogeneity of EPCAM^+^PLET1^+^ cells in the E10.5 and E12.5 thymic primordium. **(A)** Schematic showing experimental design. Thymic primordia were microdissected from E10.5 and E12.5 wild-type embryos and dissociated to single cell suspensions. Single E10.5 and E12.5 EPCAM^+^PLET1^+^ TEC were then deposited into BD Precise WTA Single Cell Encoding plates by flow cytometry processed for single cell transcriptome sequencing using BD Precise WTA reagents. **(B, C)** tSNE visualizing the clustering results with Seurat colored by cluster **(B)** and developmental stage **(C)** (n=709 cells). **(D)** Heat map showing differentially expressed genes among the eight clusters. **(E)** Expression of Notch signalling related plus selected other genes across all clusters. **(F)** Violin plot showing expression of selected markers across all clusters.

Clusters 1, 3 and 4 expressed relatively high levels of the gene encoding Placenta expressed transcript 1 (*Plet1*), a known marker of E10.5 pharyngeal endoderm (52) and one of the proteins used to select the sequenced cell population (see Materials and Methods). These clusters also expressed insulin-like growth factor binding proteins 4 (*Igfbp4*) and 5 (*Igfbp5*) and the E3 ubiquitin-protein ligase DTX4 (*Dtx4*), a known regulator of Notch signalling, with Cluster 1 also expressing high levels of metallothionein 1 (*Mt1*) and 2 (*Mt2*). Cluster 2 was characterized by expression of Neurogenic differentiation 1 (*Neurod1*) and other neural lineage markers and by low expression of Paired box 9 (*Pax9*), consistent with previous observations (36). Cluster 2 also expressed cytoskeleton-associated genes including Tubulin beta 3 (*Tubb3*), Thymosin beta 4 (*Tmsb4x*), Coactosin like F-actin binding protein 1 (*Cotl1*), alpha Tubulin (*Tuba1a*) and the neural skeleton genes Neurofilament medium chain (*Nefm*) and Neurofilament light polypeptide (*Nefl*). Clusters 1-4 were also characterized by expression of a number of genes, including the heparin-binding protein Midkine (*Mdk*), that were not expressed or expressed at only very low levels in Clusters 5-8. Of note however is that Clusters 5 and 6 expressed the Midkine family member Pleiotrophin (*Ptn*), encoding PTN which is functionally redundant with MDK.

The cells in Clusters 5 and 6 were similar and also shared similarities with Cluster 7 cells. All three clusters expressed TEC-associated genes including *Foxn1*, Paired box 1 (*Pax1*), Cytokeratin 8 (*Krt8*) and the FOXN1 targets Serine protease 16 (*Prss16*), Chemokine (C-C motif) ligand 25 (*Ccl25*) and Proteasome subunit 11 (*Psmb11*) (among others) (53). Clusters 5 and 6 also expressed *Psmb10* but did not express *Krt5.* Expression of *Krt8* and *Psmb10* was higher in Cluster 6 than in Cluster 5, suggesting Cluster 6 cells might be more differentiated than Cluster 5. Cluster 7 expressed very high levels of *Krt5, Krt17*, *Krt19, Plet1* and Claudin 4 (*Cldn4*). Thus, these data suggested that Clusters 5 and 6 might be cTEC-fated and Cluster 7 mTEC-fated; alternatively, either Cluster 5 or Cluster 7 might represent bipotent thymic epithelial progenitor cells (TEPC). Cluster 8 corresponded to parathyroid primordium cells, as evidenced by the expression of the genes encoding Parathyroid hormone (*Pth*), Chromogranin A (*Chga*) (Figure 1B-E). Examination of Notch signalling components revealed that in E12.5 TEC, *Notch1*, *Notch3* and Jagged 1 (*Jag1)* expression was largely restricted to Cluster 7 (Figure 1E), also consistent with Cluster 7 cells being mTEC or common TEC progenitors (33, 34). We note that Clusters 4, 5, 6 and 7 all contained cells that expressed *Psmb11*, despite their separate identities (Figure 1D, E), and that Cluster 7 cells expressed high levels of the genes encoding the cell surface markers Uroplakin 2 (*Upk2*) and CD9 (*Cd9*) (Figure 1E) .

Taken together these data establish that the E12.5 thymic primordium contains at least three TEC subpopulations and that the gene expression profile of Cluster 7 TEC is consistent with that of the mTEC progenitor population predicted by our previous studies, which demonstrated divergence of the mTEC and cTEC lineages prior to E11.25-12.0 (33). These findings were consistent with those of Gao (35) and Magaletta (36), who previously reported heterogeneity among TEC at E12.5 including the existence of a *Krt5^hi^* subpopulation at E12.5 enriched for *Cldn3/4*; Gao and colleagues also suggested the existence of mTEC-specified progenitor cells by E11.5 (35).

### Sox9^+^ cells are mTEC-fate restricted

Cluster 7 cells were present in the E12.5 thymic primordium at a frequency of ∼9% (Figure 1). The above data led us to consider whether these cells represented a common-or an mTEC-specific progenitor cell type. Therefore, we examined our scRNAseq data in more detail, looking in particular at transcription factor profiles (Figure 2A). Overlapping but distinct regulatory networks were associated with each of the clusters (Figure 2A, B). These data identified SRY-box transcription factor 9 (*Sox9*) and CCAAT enhancer binding protein beta (*Cebpb*) as upregulated in Cluster 7 compared to other clusters and also identified each of these transcription factors as nodes in a regulome (Figure 1F, Figure 2A, B), suggesting possible roles in the regulation of Cluster 7 cell identity or differentiation.

**Figure 2:**
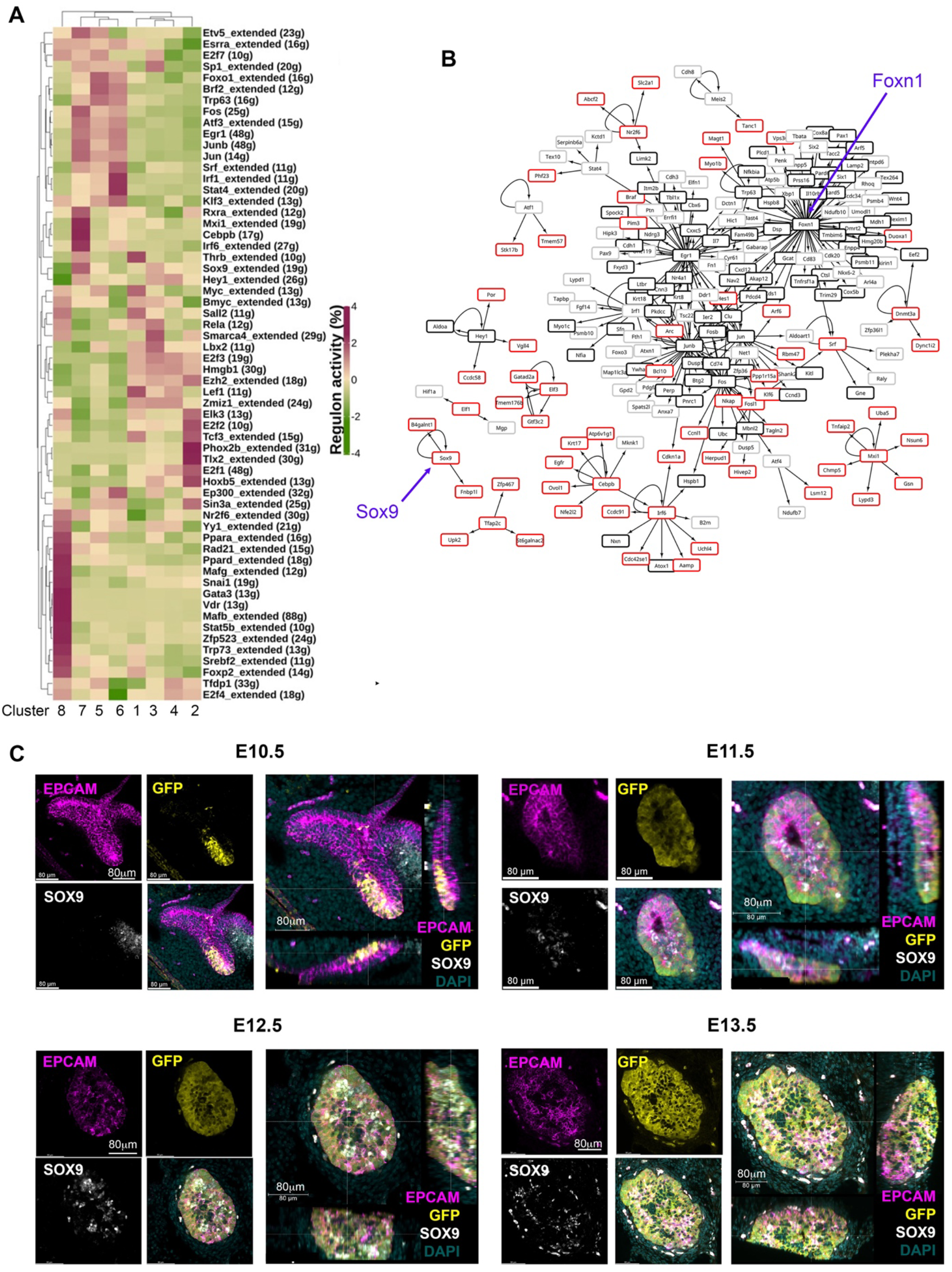
Identification of candidate regulatory networks in E10.5 and E12.5 TEC. **(A)** Heat map shows SCENIC scores of transcription factors relevant to each cluster. **(B)** Network constructed with regulons and their control targets. Arrows indicate control relationships. Red boxes show genes upregulated in Cluster 7 but not in Cluster 5 or 6. Black boxes show genes expressed in all E12.5 TEC clusters. Grey boxes show genes upregulated in E12.5 Clusters 5 and 6 compared to Cluster 7. Blue arrows highlight *Sox9* and *Foxn1* nodes. **(C)** Images show wholemount immunostaining of thick transverse sections of the pharyngeal region containing the thymic primordia from *Foxn1^GFP^* embryos at the ages and for the markers shown. Scale bars are as indicated. Note that in the E12.5 and E13.5 images the cells immediately outside the epithelial region that stain with all markers are autofluorescent.

SOX9 is a stem/progenitor cell regulator in several lineages including the intestinal epithelium (54) and is a known Notch target in early pancreatic progenitors (55, 56), but has not yet been linked to epithelial cell development in the fetal thymus. Immunohistochemical analyses revealed expression of SOX9 in the thymus domain of the 3PP from E11.5, with this staining associated with epithelial cells adjacent to the lumen of the developing primordium, and a more scattered SOX9 staining pattern at E12.5 and E13.5 (Figure 2C). No SOX9 staining was detected at E10.5 (Figure 2C). To test whether *Sox9^+^* cells in the early thymic primordium were mTEC-fated or might represent a common/bipotent TEPC, we investigated their fate *in vivo* using lineage tracing. For this, we used a model in which, in the presence of tamoxifen, CreER expressed under the *Sox9* promoter (Sox9CreER^T2^ (40)) drives activation of an inducible tdTomato reporter allele (*Gt(ROSA)26Sor^tm14(CAG-tdTomato)Hze^* [called Ai14] (41)). Briefly, Sox9CreER^T2^;Ai14 males were crossed with wild type or Sox9CreER^T2^;Ai14 females and tamoxifen was administered to pregnant females E11.5 or E13.5 (Figure 3A). The date of tamoxifen administration was taken as day 0. Embryos from mice gavaged at E13.5 were collected at day 2 (E15.5) and the thymi analyzed for the presence of tdTomato^+^ TEC. At this +2 days timepoint a very small number of tdTomato^+^ TEC, around two to three cells per lobe, were present in each of the thymi analyzed (Figure 3B), and these cells were all UEA1^+^. Thus, Sox9CreER^T2^ appeared to faithfully report SOX9 expression in the thymus (Figure 3B); only a low level of recombination was observed even at the high tamoxifen dose used, likely reflecting the relatively low expression of SOX9 in TEC at this developmental stage. We next analyzed thymi at day 5 or 7 post tamoxifen injection (i.e. E18.5, day 5 for mice treated at E13.5, day 7 for mice treated at E11.5). tdTomato^+^ TEC were present in the thymi of mice in which tamoxifen treatment had been initiated at either E11.5 or E13.5, and were exclusively located in UEA1^+^ medullary regions (shown in Figure 3C for embryos treated at E13.5 and analyzed at E18.5; note that the number of tdTom^+^UEA1^+^ cells per lobe is substantially increased at E18.5 versus E15.5). Embryos in which the tamoxifen treatment was initiated at E11.5 showed impaired development versus controls due to the high dose of tamoxifen used. However, tdTomato^+^ TEC were also restricted to UEA1^+^ regions in these thymi (not shown). No tdTomato cells were present in the negative control embryos (tamoxifen-treated Sox9CreER^T2-^ ;Ai14^+^ embryos [CreER-negative controls] and Sox9CreER^T2+^;Ai14^+^ embryos not treated with tamoxifen; Figure 3B, C). Collectively, these data identify Cluster 7 cells as mTEC-fated TEC. Furthermore, as increased numbers of tdTomato^+^ cells were observed at E18.5 versus E15.5, we conclude that *Sox9* marks mTEC-restricted progenitors.

**Figure 3:**
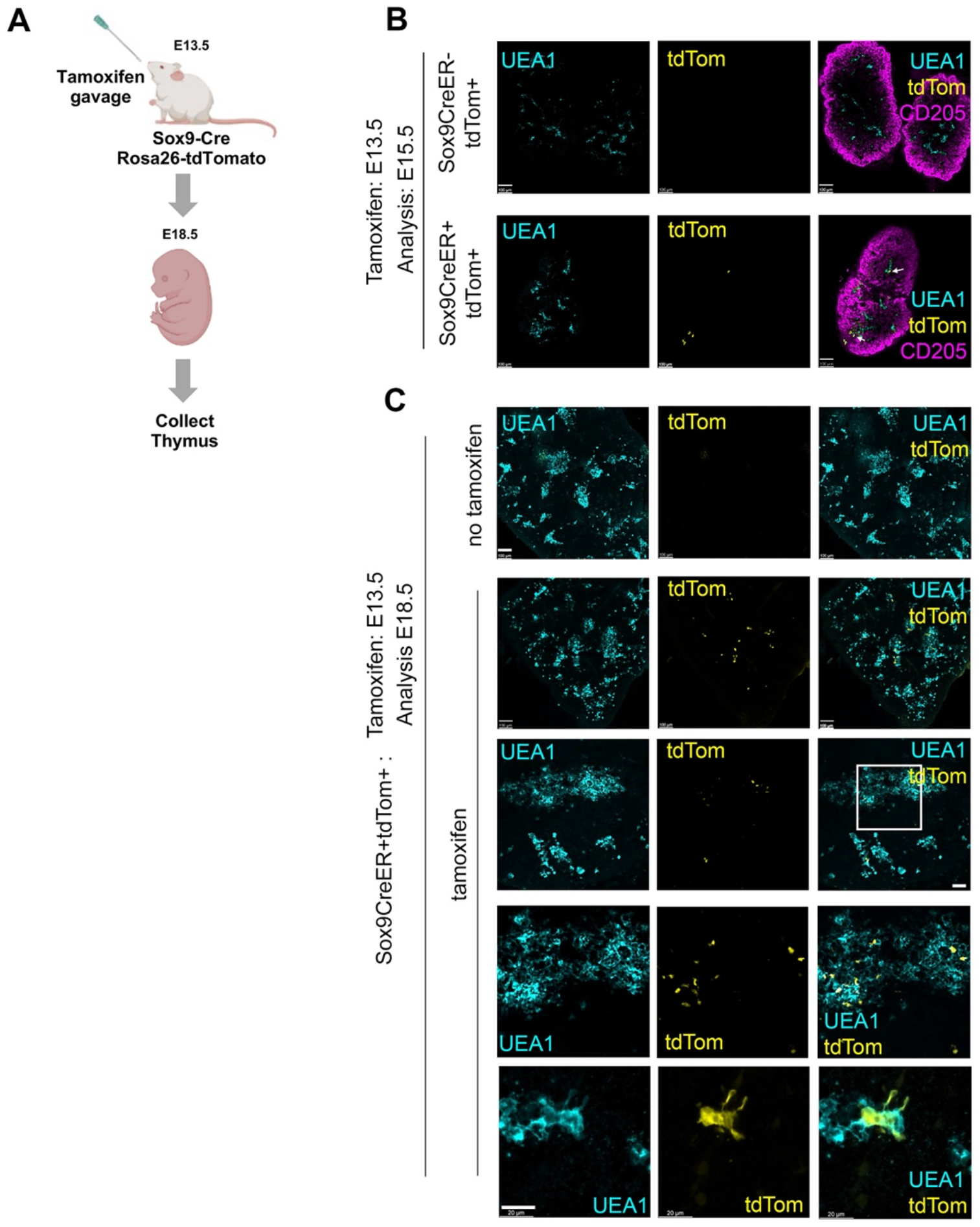
Lineage tracing demonstrates mTEC restriction of E12.5 and E13.5 Sox9^+^ TEC. **(A)** Schematic of experimental protocol. Sox9CreER;Rosa26-tdTomato (Ai14) males were crossed with wild type or Sox9CreER;Rosa26-tdTomato female mice to obtain timed matings. The pregnant females were treated with tamoxifen at E13.5. Thymi were collected at E15.5 or E18.5, genotyped and processed for wholemount imaging. **(B, C)** Images show E15.5 and E18.5 Sox9CreER^+^;Rosa26-tdTomato^+^ thymi from tamoxifen-treated and carrier-only treated embryos, after staining for the markers shown. **(B,C)** Cyan, UEA1; yellow, tdTomato; magenta, CD205. Note, in some samples penetration of αCD205 was poor. Second to bottom row in (C) shows detail from boxed area in row above. Bottom row in (C) high magnification image showing UEA1^+^tdTom^+^ cell from another section. Images show optical sections from thick sections or whole thymi. Scale bars all 100μm except C bottom row, which is 20 μm. (B) E15.5, n=3 tamoxifen-treated Sox9CreER^+^Rosa26-tdTomato^+^ embryos and n=1 tamoxifen-treated Sox9CreER^-^Rosa26-tdTomato^+^ (i.e. CreER negative control) from one pregnant female; E18.5, n=6 tamoxifen-treated Sox9CreER^+^Rosa26-tdTomato^+^ embryos from two pregnant female mice set up on different days and n=3 untreated Sox9CreER^+^Rosa26-tdTomato^+^ embryos from one pregnant female (negative controls).

### Most E12.5 TEC are cTEC-fated

The above data prompted us to revisit the potency of single epithelial cells within the early thymus rudiment, and in particular the question of whether a common TEPC exists within the E12.5 thymic primordium. For this, we used similar assay to that previously described by Anderson and colleagues (21). In brief, fetal TEC that carried a heritable genetic label were placed within an environment able to provide all the signals required for normal thymus development, then examined for the lineage contribution of their descendants after a defined time period. PLET1 is well established to mark thymic epithelial progenitor cells (TEPCs) in the E12.5 thymic primordium (51, 57). Specifically, we isolated E12.5 PLET1^+^ TEC from mice that constitutively and ubiquitously expressed a membrane-bound GFP (mGFP), by flow cytometry. The cells were then diluted to allow visual selection with a hand-controlled micropipette, and defined numbers of cells were subsequently microinjected into freshly microdissected wild-type E12.5 thymic lobes that had been trimmed of excess mesenchyme to allow easy entry of the injection pipette (Figure 4A,B). On the same day that they were microdissected and injected (within two hours after injection), the injected lobes were grafted under the kidney capsule of wild type mouse (57, 58, 59, 60, 61). The grafts were recovered after ∼14 days and examined in wholemount for mGFP^+^ cells using a fluorescence microscope (Figure 4C-E). Grafts in which mGFP^+^ cells were observed were cryosectioned and processed for immunohistochemistry. Staining for cytokeratin 14 (K14) or UEA1 (both medullary TEC markers) and CDR1 or Ly51 (both cortical TEC markers) was used to identify the location and identity of the injected mGFP^+^ cells and their progeny. DAPI was also used to distinguish cortical and medullary regions. An α-GFP antibody was used to detect injected cells. Grafts were scored by observing all recovered grafts in which mGFP^+^ foci were visible in wholemount, and analysing by immunohistochemistry every section from each graft in which mGFP^+^ foci were present.

We initially injected the maximum number of cells (sixty) that was achievable with the hand controlled micropipette. Analysis of these grafts established that mGFP^+^ cells could survive and contribute to cortical and medullary TEC networks, and to the cortico-medullary junction (CMJ). In subsequent experiments, lower defined cell numbers were injected and mGFP^+^ cells were found in a proportion of the recovered grafts. When mGFP^+^ cells were present, they typically existed as small regionalized mGFP^+^ epithelial clusters (foci) regardless of input mGFP^+^ cell number (Figure 4C, D). The results observed are shown in Figure 4 and Table 1. In many of the recovered grafts, mGFP^+^ clusters were found only in cortical regions (Figure 4C, E, F; Table 1). When sixty E12.5 PLET1^+^mGFP^+^ cells were injected into E12.5 thymic lobes and grafted under the kidney capsule, five of eight grafts analysed had mGFP^+^ cells only in the cortex (Figure 4C; Table 1) with the number of cortical foci ranging from five-to-ten to forty-to-fifty in the eight grafts analysed. One of eight grafts each showed contribution to both medulla and cortex; contribution to the CMJ and cortex; or contribution to all three compartments respectively (shown for cortical and medullary contribution in Figure 4D; Table 1). In the two grafts that had contribution to both compartments, the number of mGFP^+^ foci was greater in the cortex (Table 1). A similar picture was observed when twenty-to-thirty, ten or five E12.5 PLET1^+^mGFP^+^ TEC were injected into E12.5 thymic lobes (Figure 4E; Table 1). In none of these experiments did we recover grafts where the injected E12.5 PLET1^+^mGFP^+^ TECs injected gave rise to mGFP^+^ foci only in the medulla. Finally, we injected each E12.5 lobe with a single GFP^+^PLET1^+^ TEC: here, mGFP^+^ cells were observed in only one of eighty three grafts performed when the lobe was grafted on the same day as the injection, and were located solely in the cortex (Figure 4F; Table 1).

**Figure 4:**
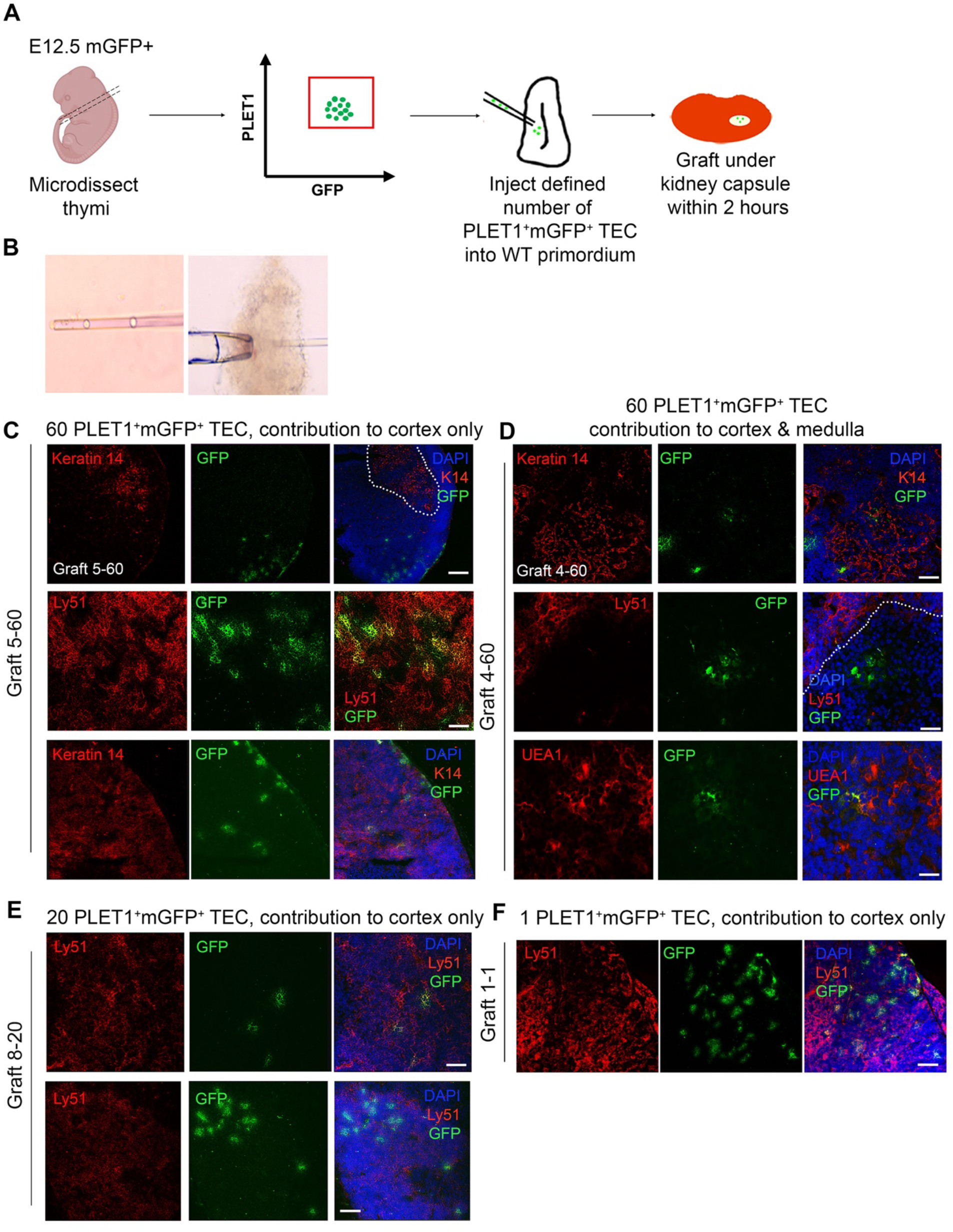
The E12.5 mGFP^+^ PLET1^+^ TEC population contains cTEC restricted and common or mTEC restricted progenitors. (A) Schematic showing experiment design. E12.5 mGFP^+^PLET1^+^ TEC were isolated from E12.5 thymi, individual cells were picked up in a hand-pulled microinjection pipette. A defined number of mGFP^+^ TEC was then injected into each wild type E12.5 thymus primordium. The injected primordia were grafted under the kidney capsule of recipient mice on the same day they were microdissected and within 2 hours of microinjection. Grafts were recovered after 2-3 weeks and analysed by histology and immunohistochemical analysis with the markers shown. (**B**) Images show individual cells within microinjection pipette (left) and E12.5 primordium held with a holding pipette whilst being injected with a microinjection pipette. **(C-F)** Images show representative staining of cortical **(C, E, F)** and medullary **(D)** mGFP^+^ foci. Sixty **(C, D),** twenty **(E)** or one **(F)** cells were injected per lobe. Cytokeratin 14 (K14) and UEA1 stain mTEC while Ly51 stains cTEC. DAPI reveals nuclei and can be used to differentiate cortical and medullary regions based on cell density. Scale bars 55μm, except C upper row, E lower row and F where scale bars represent 150μm. Graft names correspond to those in Table 1. Images show single optical sections. Dotted line in (C) top row demarcates medullary area. n for each condition represents an independent graft and is as indicated in Table 1; at least three biologically independent replicates were performed for each injection condition.

**Table 1:** E12.5 TEC preferentially adopt a cortical TEC fate. Table shows distribution of foci in grafted E12.5 thymic lobes that had been injected with 1, 5, 10, 20-30 or 60 E12.5 mGFP^+^PLET1^+^ TEC. Grafted were placed under the kidney capsule on the same day and recovered for analysis two-three weeks later. mGFP^+^ foci were scored visually for regional localization based on the DAPI, α-K14, Ly51 and CDR1 staining. Some grafts contained mGFP^+^ TEC in both the cortical and medullary compartments but most had mGFP^+^ cells in the cortex only. No grafts were scored with contribution to the medulla only. 60 cells injected, n= 8 independent grafts of which 8 grafts contained mGFP^+^ cells (n=8/8); 20-30 cells injected, n= 22 independent grafts of which 8 grafts contained mGFP^+^ cells (n=8/22); 10 cells injected, n= 8 independent grafts of which 2 grafts contained mGFP^+^ cells (n=2/8); 5 cells injected, n= 18 independent grafts of which 1 graft contained mGFP^+^ cells (n=1/18); 1 cell injected, n= 83 independent grafts of which 1 graft contained mGFP^+^ cells (n=1/83). N for each condition represents an independent graft; at least three biologically independent replicates were performed for each injection condition. Only grafts that contained mGFP^+^ cells upon visual inspection were analyzed by immunohistochemistry.

| Location and number of mGFP<sup>+</sup> foci in recovered graft |  |  |  |
| --- | --- | --- | --- |
| Graft ID | Cortex | Medulla | CMJ |
| 60 E12.5 PLET1<sup>+</sup>mGFP<sup>+</sup> cells injected into each E12.5 lobe |  |  |  |
| G1-60 | 40-45 | 0 | 0 |
| G2-60 | 10-15 | 6 | 2 |
| G3-60 | 15-20 | 0 | 3 |
| G4-60 | 5-10 | 2 | 0 |
| G5-60 | 30-35 | 0 | 0 |
| G6-60 | 15-20 | 0 | 0 |
| G7-60 | 5-10 | 0 | 0 |
| G8-60 | 5-10 | 0 | 0 |
| 20-30 E12.5 PLET1<sup>+</sup>mGFP<sup>+</sup> cells injected into each E12.5 lobe |  |  |  |
| G1-20 | 0-5 | 4 | 2 |
| G2-20 | 10-15 | 0 | 0 |
| G3-20 | 5-10 | 0 | 0 |
| G4-20 | 10-15 | 6 | 2 |
| G5-20 | 5-10 | 0 | 0 |
| G6-20 | 5-10 | 0 | 0 |
| G7-20 | 5-10 | 0 | 0 |
| G8-20 | 5-10 | 0 | 0 |
| 10 E12.5 PLET1<sup>+</sup>mGFP<sup>+</sup> cells injected into each E12.5 lobe |  |  |  |
| G1-10 | 6 | 0 | 0 |
| G2-10 | 4 | 0 | 0 |
| 5 E12.5 PLET1<sup>+</sup> mGFP<sup>+</sup> cells injected into each E12.5 lobe |  |  |  |
| G1-5 | 1-5 | 0 | 0 |
| 1 E12.5 PLET1<sup>+</sup> mGFP<sup>+</sup> cell injected into each E12.5 lobe |  |  |  |
| G1-1 to G83-1 | 40-45 | 0 | 0 |

### cTEC-fate bias among E12.5 TEC is not explained by community effect

The community effect is a developmental mechanism first described by Gurdon and colleagues whereby cells require interaction with neighbouring cells in order to express a gene that allows them to complete differentiation. If cells are pre-determined they should be able to complete differentiation even in an ectopic cellular environment (62, 63, 64). To address the possibility that a community effect might explain our results we performed two experiments. First, we attempted to bias the fate of the injected cells towards the medullary lineage by injecting them into the lumen of an E11.5 thymic primordium (Figure 5A), based on the rationale that the CLDN3/4^+^ cells lining the lumen of the 3PP were identified as candidate precursors of AIRE1^+^ mTECs (26). If there was a community effect between the injected cells and the host environment, we predicted that as the cells injected into the lumen would first contact putative mTEC progenitors they would adopt a medullary fate unless already restricted to becoming cTEC. Thirty E12.5 mGFP^+^PLET1^+^ TEC were injected into the lumen of E11.5 thymic primordia (Figure 5A; Table 2), which were then grafted and recovered after two weeks. In two out of the three grafts recovered, all of cells injected contributed only to the cortex (Table 2) while the third graft had GFP^+^ cells that contributed to both the cortex and medullary regions although most of the foci were observed in the cortex (Table 2). These data show that placing E12.5 mGFP^+^PLET1^+^ TECs into an environment likely to be enriched for precursors for medullary TECs had no apparent influence on their lineage choice.

**Figure 5:**
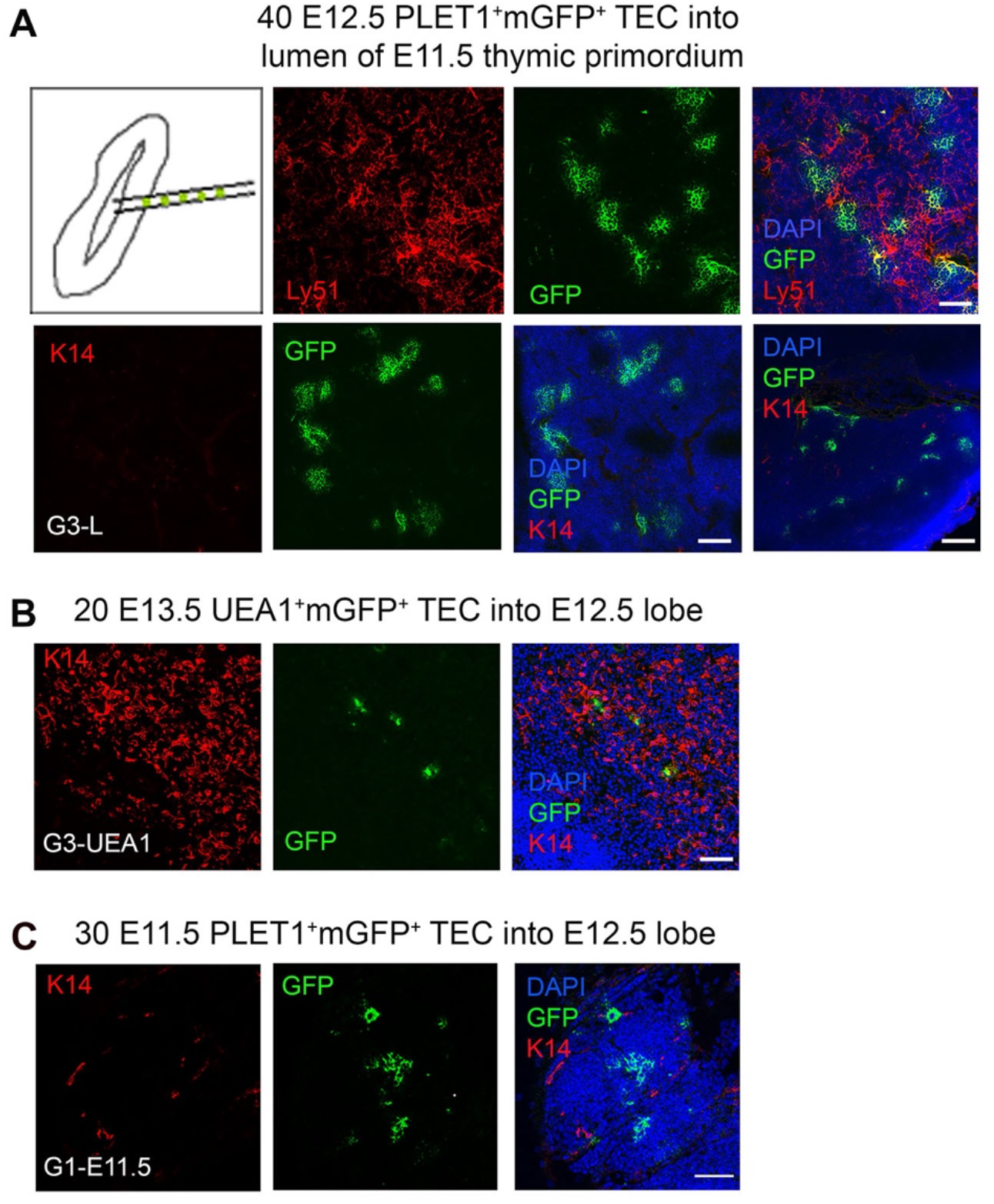
Community effect does not override cell identity in E12.5 TEC progenitors. E11.5 mGFP^+^PLET1^+^, E12.5 mGFP^+^PLET1^+^ or E13.5 UEA1^+^ TEC were isolated. A defined number of mGFP^+^ TEC was then injected into the lumen of E11.5 **(A)** or the body of wild type E12.5 **(B,C)** thymic primordia. The injected primordia were grafted under the kidney capsule of recipient mice on the same day that they were microdissected and within 2 hours of microinjection. Grafts were recovered after 2-3 weeks and analyzed by histology and immunohistochemical analysis with the markers shown. **(A)** Images show representative mGFP^+^ foci in grafts of E11.5 thymic lobes into which forty E12.5 mGFP^+^PLET1^+^ cells were injected in the lumen. mGFP^+^ foci are negative for cytokeratin 14 (K14) and positive for Ly51. **(B)** Images show representative mGFP^+^ foci in a graft in which twenty E13.5 mGFP^+^UEA1^+^ TEC were injected into the body of an E12.5 lobe. mGFP^+^ cells were located in the medulla region or the CMJ. **(C)** Images show representative mGFP^+^ foci in graft in which thirty E11.5 mGFP^+^PLET1^+^ TEC were injected into the body of an E12.5 lobe. mGFP^+^ cells were located only in the cortical region. **(A-C)** Graft names correspond to the graft identities in Table 2. Scale bars, **(A)** A 75µm except bottom right panel, 300µm. **(B)** 75µm **(C)** 55µm. N for each condition represents an independent graft and is as indicated in Table 2; at least three biologically independent replicates were performed for each injection condition.

**Table 2:**
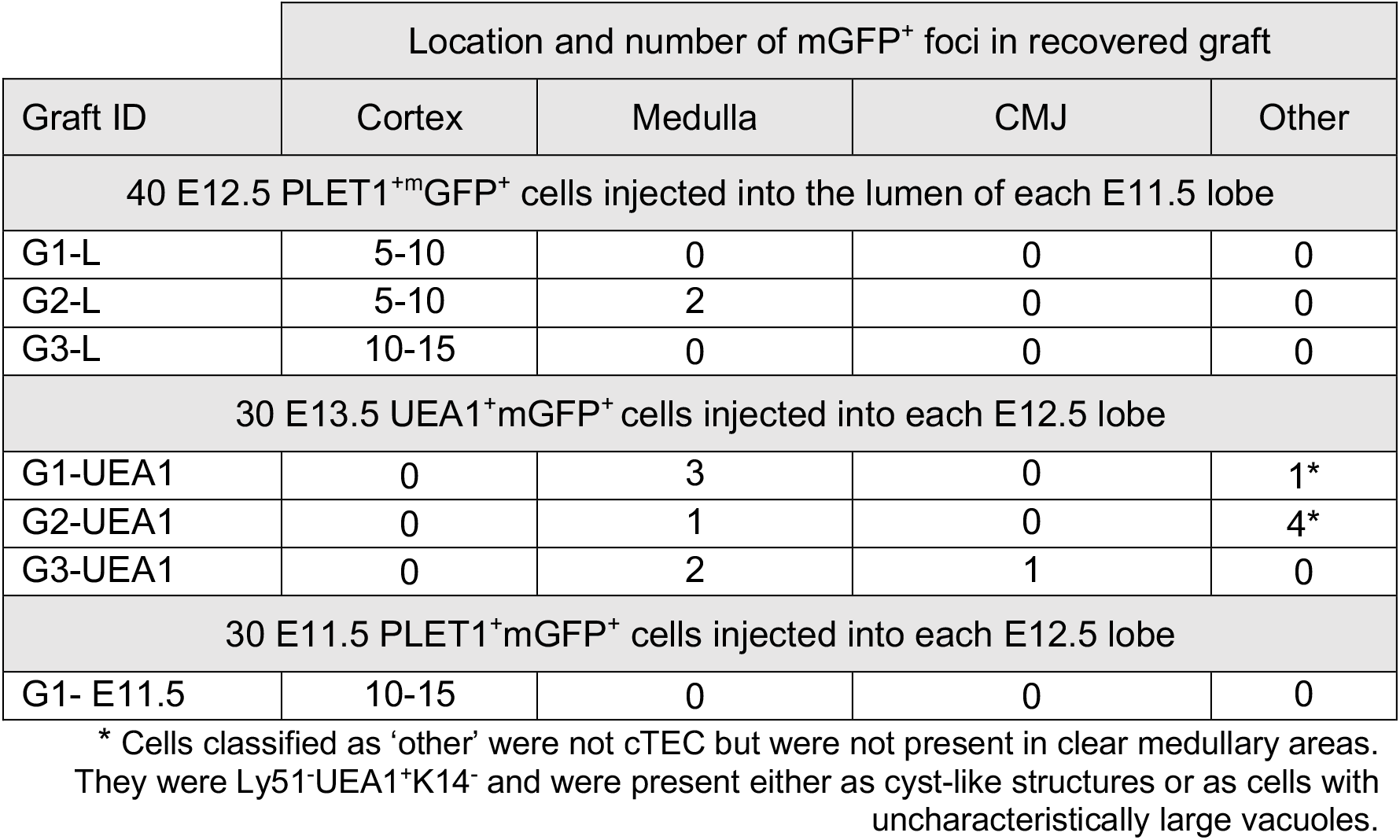
Cell identity not community effect determines progeny location. . Table shows distribution of foci in grafted E12.5 or E11.5 thymic lobes that had been injected with 20 or 40 mGFP^+^PLET1^+^ TEC of the developmental age shown. The injected cells were placed in either the body or the lumen of the lobe, as indicated. Grafted were placed under the kidney capsule on the same day and recovered for analysis two-three weeks later. mGFP^+^ foci were scored visually for regional localization based on the DAPI, α-K14, Ly51 and CDR1 staining. 40 E12.5 mGFP^+^PLET1^+^ TEC into E11.5 lobe lumen, n≥5 independent grafts of which 3 grafts contained mGFP^+^ cells (n=3/≥5); 30 E13.5 mGFP^+^UEA1^+^ TEC injected into E12.5 lobe body, n= 40 independent grafts of which 3 grafts contained mGFP^+^ cells (n=3/40); 30 E11.5 mGFP^+^PLET1^+^ TEC injected E12.5 lobe body, n= 4 independent grafts of which 1 graft contained mGFP^+^ cells (n=1/4). N for each condition represents an independent graft; at least three biologically independent replicates were performed for each injection condition. Only grafts that contained mGFP^+^ cells upon visual inspection were analyzed by immunohistochemistry.

Second, Hamazaki and colleagues previously showed that CLDN3/4^hi^UEA1^+^ TEC, when reaggregated with dissociated E13.5 whole thymic lobes, gave rise to medullary but not cortical TECs (26). Therefore, we isolated E13.5 mGFP^+^UEA1^+^ TEC and injected 30 cells into each E12.5 thymic lobe before grafting under the kidney capsule (Figure 5B). The grafts were recovered after three weeks and the fate of the mGFP^+^UEA1^+^ cells was assessed. In these analyses, mGFP^+^ foci were most frequently found in K14^+^ medullary regions (Figure 5B; Table 2). Some mGFP^+^ cells were observed in the cortical regions, but these were in cyst like structures or did not appear to be cortical cells as they had uncharacteristically large vacuoles. These cortical GFP^+^ structures were positive for UEA1 but not for K14 or Ly51. Thus, medullary-restricted UEA1^+^mGFP^+^ TEC did not become cTEC in our assay.

Collectively, these data strongly suggest that when mGFP^+^PLET1^+^ TEC are injected into a normal fetal thymic microenvironment this does not override their existing lineage-identity, providing further support for the existence of sub-lineage precursors for the cortical and medullary compartments by E12.5. To test whether this might also be the case at earlier developmental stages, we also injected thirty E11.5 mGFP^+^PLET1^+^ cells into E12.5 thymic lobes. In these experiments, mGFP^+^ clusters were found only in the cortex (shown in Figure 5C by lack of K14 staining and dense DAPI staining) suggesting that as early as E11.5 most TEC are committed to a cortical fate (Figure 5C; Table 2), consistent with a recent scRNAseq analysis (36).

Taken together, the data presented in Figures 1-5 strongly suggest that at E12.5 the majority of TEC are already specified to the cortical fate and further demonstrate that at this developmental stage, a minor population is mTEC-fated. These findings are consistent with the picture emerging from scRNAseq and barcoding analyses (35, 36, 37) but contrast with the conclusions of Rossi and colleagues (21). This raised the obvious question of why our data differed from those of Rossi (21) who showed via a very similar microinjecting method that one injected E12.5 YFP^+^ TEPC could give rise to cells scattered evenly throughout both the cortex and medulla, concluding the existence of bi-potent TEPC at E12.5. This result was observed in all four grafts in which YFP^+^ cells were found, of a total of thirteen grafts injected (21).

Rossi and colleagues used an α-EPCAM rather than α-PLET1 antibody to isolate the YFP^+^ TEC, and showed that these antibodies both bind most if not all TEC in the E12.5 population (21). Therefore, we repeated our analyses substituting α-EPCAM for α-PLET1 as the antibody used to isolate the TEC for injection. Analysis of these grafts gave identical results to those shown above (Supplementary Figure 3). We considered the potential effect of the method used to dissociate the fetal thymic lobes in the two laboratories. In the analyses presented above, cells were dissociated using collagenase and hyaluronidase, whereas Rossi and colleagues used trypsin/EDTA for 5 minutes at 37°C (21). Thus, we also tested the outcome of injecting twenty-to-thirty cells obtained by after dissociation in trypsin/EDTA. No difference in outcome was observed (data not shown). We also ruled out selective immune rejection: in the above experiments, we sorted cells from lobes collected from a C57BL/6 x mGFP^+^C57BL/6 cross and then grafted into C57BL/6xCBA F1 hosts. Although the F1 host should not reject cells from either parental strain, we tested the outcome of injecting E12.5 TEC from a mGFP^+^C57BL/6 x CBA cross. Again, the same results were observed (Supplementary Figure 4). Furthermore, use of an αGFP antibody gave the same results as direct detection of mGFP (Supplementary Figure 4).

### Overnight culture changes the potency of E12.5 TEC

It is well established that *in vitro* culture in incompletely defined culture medium can affect cell potency. Therefore, we tested the effect of altering our protocol such that rather than grafting the lobes within two hours of injection, we cultured the injected lobes overnight in DMEM 10% FCS prior to grafting. When these grafts were analyzed, four of twenty two contained mGFP^+^ foci and in all four, these were found in both cortical and medullary regions (Figure 6; Table 3). In one of the grafts, most of the mGFP^+^ cells were found in the medulla and expressed Keratin 14 (Figure 6B). The mGFP^+^ clusters found in cortical regions expressed CDR1, a marker restricted to cortical TEC of the adult thymus (Figure 6B). Overnight culture of either the E12.5 mGFP^+^PLET1^+^ TEC or the injected lobes prior to grafting also increased the frequency of grafts containing medullary foci (Tables 3 and 4, Supplementary Figure 5). When thirty-to-forty E12.5 mGFP^+^PLET1^+^ were cultured overnight then injected into E12.5 lobes and grafted within two hours, three of three grafts showed a contribution to both cortical and medullary compartments (Figure 6; Table 3). Similarly, when E12.5 lobes were cultured overnight then injected with thirty to forty mGFP^+^PLET1^+^ E12.5 TEC, three of eighteen grafts showed a contribution to both compartments (Tables 3 and 4; Supplementary Figure 5).

**Figure 6:**
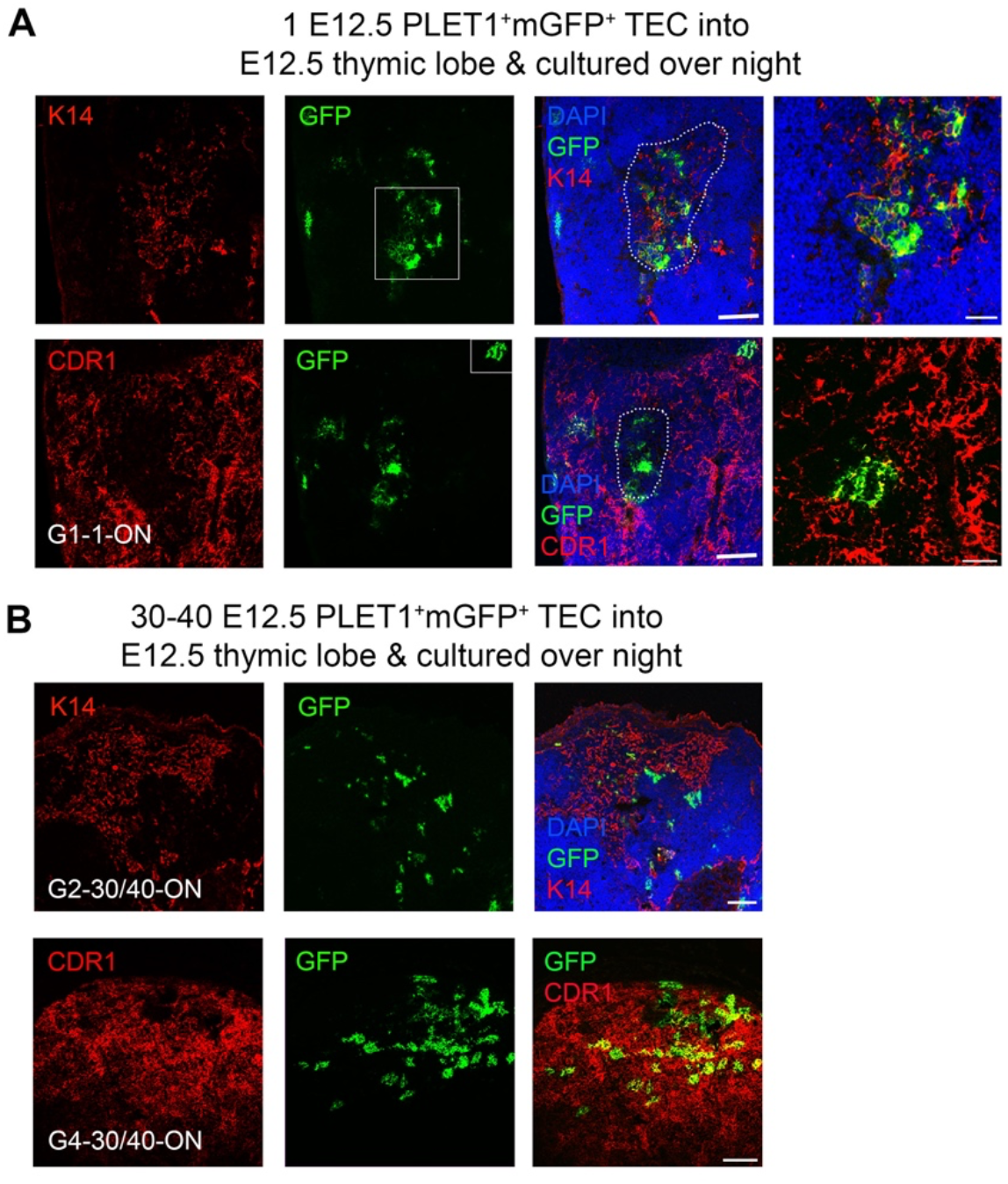
A bipotent thymic epithelial progenitor cell is present after overnight culture. E12.5 mGFP^+^PLET1^+^ TEC were isolated from E12.5 thymi and a single cell was then injected into each wild type E12.5 thymus primordium; injected primordia were cultured overnight before grafting under the kidney capsule of recipient mice, grafts were recovered after 2-3 weeks and analysed with the markers shown. **(A)** Representative mGFP^+^ foci derived from a single cell showing co-localisation of mGFP^+^ cells with the mTEC marker Keratin 14 (k14) and the cTEC marker CDR1. DAPI shows nuclei. Right hand panels show higher magnification of boxed regions. **(B)** Representative mGFP^+^ foci from grafts of E12.5 thymic lobes injected with 40 mGFP^+^Plet1^+^ TEC and then cultured for 24 hours before grafting. Images show immunostaining for anti-K14 and CDR1 in two separate grafts. Graft name corresponds with graft identification in Table 3. Scale bars **(A)** 150μm except right hand panels, 55μm, **(B)** 150μm. N for each condition represents an independent graft and is as indicated in Table 3; at least three biologically independent replicates were performed for each injection condition.

**Table 3:** Potency of E12.5 TEC is changed by overnight culture. Table shows distribution of foci in grafted E12.5 thymic lobes that had been injected with the number of E12.5 mGFP^+^PLET1^+^ TEC shown. The injected lobes were cultured overnight before grafting under the kidney capsule. Grafts were recovered for analysis after two-three weeks. mGFP^+^ foci were scored visually for regional localization based on the DAPI, α-K14, Ly51 and CDR1 staining. 30-40 E12.5 mGFP^+^PLET1^+^ TEC injected into E12.5 lobe then cultured, n=4 independent grafts of which 4 grafts contained mGFP^+^ cells (n=4/4); 30-40 E12.5 mGFP^+^PLET1^+^ TEC cultured then injected into E12.5 lobe, n≥5 independent grafts of which 3 grafts contained mGFP^+^ cells, (n=3/≥5); E12.5 lobes cultured overnight then injected with 30-40 E12.5 mGFP^+^PLET1^+^ TEC, n=18 independent grafts of which 3 grafts contained mGFP^+^ cells (n=3/18); 1 E12.5 mGFP^+^PLET1^+^ TEC injected into E12.5 lobe, n=22 independent grafts of which 4 grafts contained mGFP^+^ cells (n=4/22). Only recovered grafts that contained mGFP^+^ TEC upon visual inspection were analyzed by immunohistochemistry. Contribution to the medulla appeared greater in grafts that were cultured overnight (O/N) although one graft was observed with contribution to only the cortex. N for each condition represents an independent graft; at least three biologically independent replicates were performed for each injection condition. The proportion of cTEC-fated mGFP^+^ cells in grafted lobes that had not been (and in which no cells had been) cultured (Table 1) with that in grafted lobes that had been cultured overnight before grafting (Table 3) was statistically significant as shown by a pooled two-proportion z-test, which gave a z-score of z=3.03 and ap-value of p-val=0.002.

| Location and number of mGFP<sup>+</sup> foci in recovered graft |  |  |  |  |
| --- | --- | --- | --- | --- |
| Graft ID | Cortex | Medulla | CMJ | Other |
| 30-40 E12.5 PLET1<sup>+</sup>mGFP<sup>+</sup> cells injected into each E12.5 lobe and cultured O/N before grafting |  |  |  |  |
| G1-30/40-O/N | 15-20 | 3 | 3 |  |
| G2-30/40-O/N | 10-15 | 10-15 | 0 |  |
| G3-30/40-O/N | 20-30 | 0 | 0 |  |
| G4-30/40-O/N | 15-20 | 5 | 0 |  |
| 30-40 E12.5 PLET1<sup>+</sup>mGFP<sup>+</sup> cells cultured overnight then injected into uncultured E12.5 lobes and grafted within 2 hours |  |  |  |  |
| G2 | 15-20 | 1-5 | 1-5 |  |
| G3 | 1-5 | 1-5 | 0 |  |
| G4 | 35-40 | 5-10 | 0 |  |
| E12.5 lobes cultured overnight then injected with 30-40 E12.5 PLET1<sup>+</sup>mGFP<sup>+</sup> cells and grafted within 2 hours |  |  |  |  |
| G1 | 1-5 | 5-10 | 1-5 |  |
| G2 | 15-20 | 1-5 | 0 |  |
| G3 | 1-5 | 1-5 | 1-5 |  |
| 1 E12.5 PLET1<sup>+</sup>mGFP<sup>+</sup> cell injected into each E12.5 lobe and cultured O/N before grafting |  |  |  |  |
| G1-1-ON | 3 | 1* | 0 |  |
| G2-1-ON | 1-5 | 1-5 | 0 |  |
| G3-1-ON | 1-5 | 1-5 | 0 |  |
| G4-1-ON | 1-5 | 1-5 | 0 |  |

\*Very large medullary focus

**Table 4.** Summary of cell potency assay data. Proportion of grafts with contribution from PLET1^+^mGFP^+^ cells, for each of the conditions shown. *E12.5 TEC injected into lumen of E11.5 lobe, **2 grafts also contained atypical GFP^+^ TEC.

| Number of injected mGFP<sup>+</sup>PLET1<sup>+</sup> cells | Number of grafts with at least one mGFP<sup>+</sup> focus/Total number of grafts performed | Percentage of grafts with at least one mGFP<sup>+</sup> focus | Percentage of mGFP<sup>+</sup> grafts with only cTEC foci | Percentage of mGFP<sup>+</sup> grafts with cTEC and CMJ and/or mTEC foci | Percentage of mGFP<sup>+</sup> grafts with only mTEC or mTEC and CMJ foci |
| --- | --- | --- | --- | --- | --- |
| **E12.5 PLET1<sup>+</sup>mGFP<sup>+</sup> TEC** |  |  |  |  |  |
| *Injection on day of microdissection, followed by grafting within 2 hours* |  |  |  |  |  |
| 60 | 8/8 | 100 | 62.5 | 37.5 | 0 |
| 40* | 3/≥5 | 100 | 67 | 33 | 0 |
| 20 | 8/22 | 36 | 75 | 25 | 0 |
| 10 | 2/8 | 25 | 100 | 0 | 0 |
| 5 | 1/18 | 5 | 100 | 0 | 0 |
| 1 | 1/83 | 1.2 | 100 | 0 | 0 |
| *Injection on day of microdissection, followed by overnight culture before grafting* |  |  |  |  |  |
| 30-40 | 4/4 | 100 | 25 | 75 | 0 |
| 1 | 4/22 | 18 | 0 | 100 | 0 |
| *Cells cultured overnight prior to injection, followed by grafting within 2 hours* |  |  |  |  |  |
| 30-40 | 3/≥5 | 100 | 0 | 100 | 0 |
| *Lobes cultured overnight prior to injection with freshly isolated cells, followed by grafting within 2 hours* |  |  |  |  |  |
| 30-40 | 3/18 | 100 | 0 | 100 | 0 |
| **E11.5 PLET1<sup>+</sup>mGFP<sup>+</sup> TEC** |  |  |  |  |  |
| *Injection on day of microdissection, followed by grafting within 2 hours* |  |  |  |  |  |
| 40 | 1/4 | 100 | 100 | 0 | 0 |
| **E13.5 UEA1<sup>+</sup>mGFP<sup>+</sup> TEC** |  |  |  |  |  |
| *Injection on day of microdissection, followed by grafting within 2 hours* |  |  |  |  |  |
| 20 | 3/40 | 100 | 0 | 0 | 100** |

Collectively, in our study a clonal result was achieved in five out of one hundred and five single cell injections (1/83 without overnight culture and 4/22 after overnight culture) (Table 4). When freshly isolated cells were injected and the lobes grafted on the same day, the single cell injected generated only cTEC progeny. In contrast, when the injected lobes were cultured overnight before grafting, each of the single injected cells that contributed to the graft was bipotent. As the frequency of single cell contribution without overnight culture was consistent with the estimated viability of mGFP^+^ cells in our experiments (Table 4) it remains possible that, as well as the minor population of mTEC-restricted progenitors demonstrated through lineage tracing (Figure 3), a common progenitor able to generate both cTEC and mTEC exists at low frequency in E11.5 and E12.5 thymic primordia. However our data strongly support a model where most TEC in the E12.5 thymus are cTEC-fated progenitors, in contrast to the data presented by Rossi and colleagues (21) and consistent with our own (Figure 1) and previously reported scRNAseq data (35, 36) and with genetic analyses reported by ourselves and others (33, 34). Furthermore, they indicate that the potency of E12.5 TEC can be changed by overnight culture.

### Molecular basis of changed potency of E12.5 thymic epithelial cells after overnight culture

To investigate the possible mechanisms responsible for the predominant bipotent TEPC activity observed after overnight culturing, we performed scRNAseq on E12.5 TEC, E13.5 TEC and TEC that were isolated at E12.5 and cultured overnight as either fetal thymic organ culture (FTOC) or monolayers, which we will refer to collectively as ON TEC (Figure 7A, B). As expected, across the whole analysis, the majority of TEC were cTEC-like (similar to Clusters 5 and 6, Figure 1) with the rest of the cells being mTEC-like (similar to Cluster 7, Figure 1). The cTEC-like cells split into two clusters, “cTEC” and “ON cTEC”, with the latter consisting almost exclusively of cells from the overnight culture condition (Figure 7A). The mTEC cells from E13.5 and ON TEC included some Aire^+^Tnfrsf11a^+^ mTEC, which were absent at E12.5 (Figure 7C).

**Figure 7:**
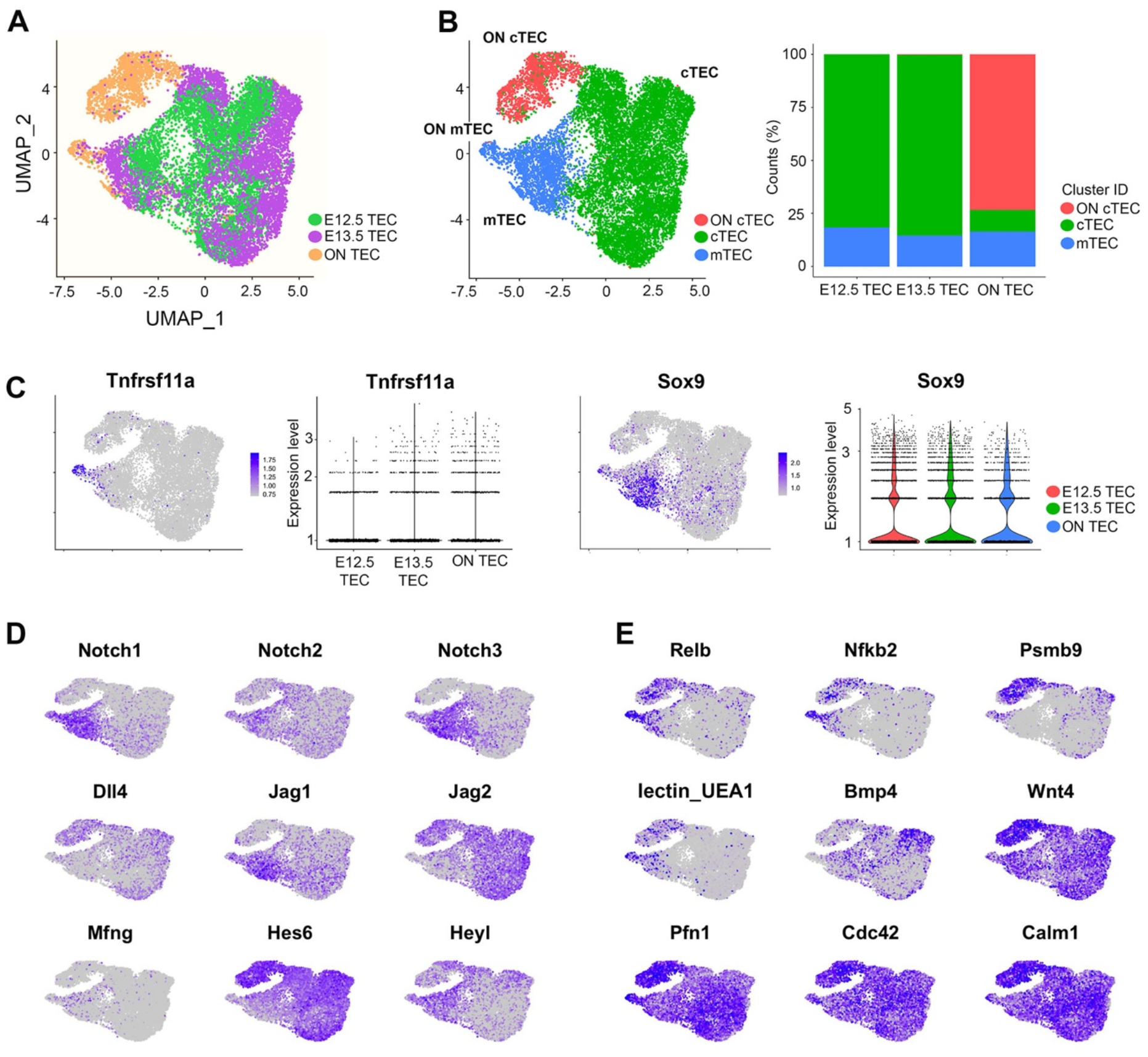
Single-cell transcriptomics reveal pathways active in overnight culture TEC (ON TEC). **(A,B)** UMAP plot of E12.5, E13.5 and ON TEC with cells colored by time point **(A)** or by cluster, with percentages of cells belonging to each cluster **(B)**. **(C)** Expression levels of the mTEC marker *Tnfrsf11a* and the early mTEPC marker *Sox9* are similar in ON TEC and E13.5 TEC. **(D)** Notch signaling genes have similar expression in ON TEC. **(E)** The canonical NF-κB targets, *Relb*, *Nfkb2* and *Psmb9*, are higher in ON TEC (top row). High UEA1 fluorescence (lectin_UEA1) suggesting a mixed phenotype (middle row). Wnt target genes, *Pfn1*, *Cdc42* and *Calm1*, are expressed more highly in ON TEC (bottom row and Supplementary Tables 4 and 5).

Since ON culture had revealed a bipotent TEPC activity not detected in the absence of culturing (Figure 5), we first checked if ON TEC contained a higher percentage of mTEC, *Sox9^+^* cells or *Tnfrsf11a^+^* cells and found no major differences (Figure 7B, C). This suggests that ON culture does not directly convert cTEC to mTEC, at least in the 24-hour period, but rather ‘reprogrammes’ the predominant cTEC-fated population to a bipotent state able to adopt both cTEC and mTEC fates. In the UMAP projection (Figure 7), the ON cTEC are closer to E13.5 cTEC than E12.5 cTEC, suggesting that the ON cTEC are not a dedifferentiated cTEC population and thus that dedifferentiation does not explain the gain in bipotency observed. Since Notch signalling is required for the mTEC fate, it was possible that changes in Notch signalling competence might be implicated. However, we found no major differences in Notch pathway genes between cTEC and ON cTEC (Figure 7D).

We then performed a differential gene expression and GO enrichment analysis comparing cTEC to ON cTEC and found evidence for high NF-κB signalling in the ON cTEC. In particular, the canonical NF-κB target genes *Relb, Nfkb2* and *Psmb9* are high in ON cTEC but very low/off in cTEC (Figure 7E and Supplementary Table 4). Since RELB is an effector of the non-canonical NF-κB pathway, this is an example of canonical NF-κB signalling priming cells for non-canonical NF-κB signalling. We note that RELB is essential for formation of the thymic medulla; *Relb^-/-^* mice thymi lack medullary compartments (28, 65, 66). The ligand responsible for induction of NF-κB signalling in the cultured cells is unknown, however, factors present in FCS are possible sources and it is also possible that factors secreted by thymocytes the ON culture condition may also contribute. For instance, some of these thymocytes express TNF (Supplementary Fig. 6; filtered out in Figure 7). The ON cTEC also showed some evidence of a mixed cTEC/mTEC phenotype, exhibiting some UEA1 fluorescence as well as higher expression of *Fut1*, which synthesises the glycan target of the UEA1 lectin (67, 68) (Fig. 7E). The ON cTEC also had lower expression of *Bmp4, Bmpr1,2* and higher expression of *Wnt4* than E12.5 or E13.5 cTEC (Fig. 7E and Supplementary Table 5). There was also evidence for higher canonical and non-canonical WNT signalling in the ON cTEC versus cTEC (Supplementary Table 4). Notably, increasing WNT signalling by knocking down the Wnt antagonist *Kremen1* was previously shown to disrupt thymus patterning (69).

From these data we conclude that overnight culture of E12.5 TEC, either as intact lobes or monolayers, changes the gene expression profile of cTEC such that it differs from that of cTEC in the E12.5 or E13.5 thymus. In particular, with the upregulation of NF-κB signalling and acquisition of a mixed phenotype with individual cells expressing genes characteristic of cTEC and of mTEC progenitors, our data suggest that the culture conditions induce a reprogramming (or possibly dedifferentiation) response that renders the cultured cTEC able to respond to signals to elaborate the mTEC-as well as the cTEC-gene expression programmes.

## Discussion

We have shown that the E12.5 thymic primordium contains a major cTEC-restricted and minor mTEC-restricted TEPC population, rather than being predominantly composed of common (bipotent) TEPC. This conclusion is supported by three lines of evidence. First, separate populations with transcriptional profiles consistent with (i) a cTEC or common TEPC and (ii) an mTEC or common TEPC identity (characterized by expression of *Sox9* and very high levels of *Krt5*) are present in the E12.5 thymus, along with (iii) a more differentiated cTEC population. Second, lineage tracing from the mTEC/common TEPC using *Sox9CreER* demonstrates the E12.5 *Krt5^very^ ^high^Sox9^+^* population we found is mTEC-fate restricted at least until E18.5, establishing this population as mTEC-restricted not common TEPC. Third, single cell transplantation experiments show that most E12.5 (and E11.5) TEC are cTEC-fate restricted, but following overnight culture most E12.5 TEC exhibit common TEPC activity reflected by acquisition of a mixed phenotype. Our data collectively support a model in which cTEC-mTEC lineage divergence in thymus organogenesis first occurs prior to E11.25, and provide functional evidence for the existence of cTEC-and mTEC-restricted progenitors prior to E12.5. We found no evidence for a predominant common TEPC population at E12.5, and indeed could only reveal this activity following overnight culture. These findings raise several points, which are discussed below.

### Functional evidence supports the presence of cTEC and mTEC restricted progenitors in the E12.5 thymus

Our data show that some TEPCs are committed to a cortical fate as early as E11.5 and that at E12.5 the majority of cells are cTEC fate-restricted. For example, when grafting 60 cells, we observed that 5/8ths of the grafts had only cortex contributions. If we denote by *N* the number of donor cells that survive the transplantation process (and are observed) and assume that each cell has an independent probability, *p*, of giving rise to just cTEC, then that would imply *p^N^*=5/8, which suggests a large proportion of cells are strictly cTEC fated. For example, if *N*=10 then *p* would be approximately 97%. Our scRNAseq shows the ratio of cTEC to mTEPC to be approximately 9:1. If *p*=0.9, then *p^N^*=5/8 would imply *N*≈4 cells, which is biologically plausible. The assumption of independent probabilities is likely inaccurate due to the spatial patterning of the thymus already present at E12.5, but even in the extreme case of completely dependent probabilities, our results still indicate that at least 5/8ths of cells are fate-restricted to the cTEC lineage (95% C.I. of 56-92% for a spatially patterned model and 95% C.I. of 73-99% for a well-mixed model with four out of 60 surviving cells, see methods). Thus, by E12.5 committed cortical lineage-restricted progenitors are present and exist in larger numbers than cells restricted to a medullary fate.

Consistent with this, the scRNAseq data presented herein (Figures 1 and 7) and by others (35, 36), show two clear TEC populations at E12.5: a large putative cTEC population (Figure 1, Clusters 5 and 6) and a smaller putative mTEPC group (Figure 1, Cluster 7). The data discussed above establish the existence of cTEC lineage restricted progenitors, presumably corresponding to Cluster 5 and 6 TEC. The putative mTEPCs (Cluster 7) express markers *Cldn3,4, Plet1, Sox9* and *Krt19* and show evidence of Notch signalling in addition to the expression of *Sox9*. This identity is consistent with that of previously identified candidate mTEPC present at E12.5, which are formed and maintained by Notch signalling and are required for medulla formation at least in the fetal thymus (26, 33, 34). Since the possibility that the Cluster 7 population represented either a common TEPC or mTEC-restricted progenitor remained open (indeed our previous studies provided some evidence for a link between Notch signalling and a common TEPC cell state (33)), we addressed these two alternatives by lineage tracing using *Sox9CreER*. Our data showed no contribution of the progeny of *Sox9^+^* cells to the cTEC lineage by E18.5, establishing the *Krt5^veryhigh^Sox9^+^Cldn3,4^+^Plet1^+^*population as mTEPC. These data are consistent with the findings of Rodewald and colleagues (25), who concluded that in mice the thymic medulla originates from around 900 mTEC-restricted progenitor cells per lobe and that these progenitors are present from E13.5. They are also consistent with the findings of Nusser and colleagues, who using *in vivo* barcoding revealed cTEC-restricted but not a common TEPC activity in the early fetal thymus (37). They further extend the findings of Gao, Magaletta and Hamazaki, who respectively identified putative mTEPC at E11.5/12.5 and identified functionally validated mTEPC at E13.5 (26, 27, 35, 36). The E12.5 *Krt5^very^ ^high^Sox9^+^Cldn3,4^+^Plet1^+^* population does not express *Tnfsfr11a* (RANK)(see Figure 1), placing it upstream of the RANK^+^ mTEPC and strongly suggesting it represents the CLDN3/4^+^SSEA1^+^ mTEPC identified by Hamazaki and colleagues (27, 28). *Sox9* is known to be regulated by Notch, also consistent with this population being the Notch-dependent very early mTEPC population recently identified by ourselves and others (33, 34). At E12.5, *Sox9^+^* (Cluster 7) cells co-express *Krt19*, which has very recently been identified as marking mTEC progenitors (70). Cluster 7 cells also specifically expressed Uroplakin 2 (*Upk2*) and *Cd9*, which encode cell surface proteins; CD9 was also identified by Lucas and colleagues as a surface marker of the *Krt19^+^* mTEC progenitor population. We note that cells in Clusters 5, 6 and 7 all express the FOXN1 target *Psmb11*. Lineage tracing using *Psmb11-Cre* (*19*), although not performed at single cell resolution or under temporal control, has been used to argue that in the fetal thymus a common cTEC-like progenitor gives rise to all TEC sub-lineage cells. β5t, the *Psmb11* gene product, is first detected in TEC at E12-12.5 (71), while eGFP activated by *Psmb11-Cre* is detected first at E12.75 (19). The finding that *Psmb11* is expressed in each of the three subpopulations of the E12.5 thymus thus reconciles the observation that all TEC arise from *Psmb11^+^* progenitors (19) with our findings that most TEC are sub-lineage restricted by E12.5.

Our finding, that most TEC at E11.5 and E12.5 are cTEC-fate restricted, contrasts with that of Rossi and colleagues who showed that most E12.5 were common TEPC (21). In trying to understand this difference, we sequentially ruled out a number of trivial explanations including assay bias resulting from a community effect at the injection site, immune rejection and minor differences in our experimental protocols: none of these accounted for the differences observed. However, we were able to replicate Rossi’s finding of a common (bipotent) TEPC when we cultured the injected lobes overnight in serum prior to transplantation under the kidney capsule. Under these conditions, we observed that most E12.5 TEC were bipotent. This finding may therefore reconcile our results with those of Rossi and colleagues (21). The change in potency in E12.5 TEC elicited by overnight culture is discussed in further detail below.

### Cell growth in the thymic primordium

In our study the descendants of the mGFP^+^ test cells, whether cTEC or mTEC, were distributed in small clusters (foci) regionalized within the recovered graft. This is consistent with the findings of Rodewald (25) and of Bleul and colleagues who, using a different lineage-tracing model, observed clustered progeny in both cortex and medulla following clonal labelling at day 14 postnatal (18).

By considering the work of Nicolas and colleagues (72, 73, 74), on the central nerve system and myotome development using retrospective clonal analysis using the LaacZ labelling system, we can begin to predict possible growth models for the thymic primordium. Nicolas defined several different modes of proliferation, dispersal and differentiation of cells *in situ* (72, 73, 74, 75). The distribution of mGFP^+^ cells in the recovered grafts would be consistent with either of two possible growth models. The mGFP^+^ cells may have remained at the site of injection, followed by a period of coherent growth in which the cells remained close to one another after each division and formed clusters. Alternatively, the mGFP^+^ cells may have undergone ordered intermingling with neighbouring cells after an initial period of proliferation, before undergoing coherent growth resulting in clusters. Since the number of foci in some of the grafts is higher than predicted if the cell viability is 1:40-1:100, the latter growth model appears to best apply.

### Broadening of fate options induced by *in vitro* culture

As discussed above, our data indicate that overnight culture in serum changes the potency of E12.5 TEC such that, rather than being cTEC-restricted, the majority acquire common (bipotent) TEPC activity. We also saw an increase in mTEC contribution in our grafting experiments when either just the recipient lobes, or just the mGFP^+^ test cell(s), were cultured overnight. This suggests that overnight culture induces the common TEPC activity through the direct action of serum components and/or by inducing a feedback loop whereby cells secrete factors that then induce the common TEPC activity in other cells. Our scRNAseq analysis identifies upregulation of NF-κB signalling activity, and acquisition of a mixed phenotype with features of both cTEC and mTEC, in cultured cTEC as possible explanations for the acquisition of bipotency in these normally cTEC-fated cells. This is reminiscent of the intestinal epithelium, where an inflammatory environment triggers the conversion of differentiated cells into stem cells through an NF-κB-WNT axis (76). Specifically, we observed an increase in canonical NF-κB signalling, including higher expression of *Relb*, which is involved in non-canonical NF-κB signalling. RELB is essential for medulla formation and NF-κB signalling plays a central role in the mTEC differentiation hierarchy (31, 65, 66, 77, 78, 79, 80). The ectopic upregulation of *RelB* and NF-κB signalling may therefore underpin the acquisition of bipotency by the cultured cTEC. In addition to factors present in serum, other potential sources of activating signal for canonical NF-κB signalling are the stress response induced by hypoxia in culture, and TNF, secreted by cultured thymocytes. A link between hypoxia and NF-κB signalling has been demonstrated, and hypoxia has been reported to affect other aspects of TEC identity (81, 82). In this regard, it is possible that similar signals modulate the potency of TEPC as thymus development progresses, consistent with the temporal changes in the balance of cTEC/mTEC contribution of individual progenitor cell clones described by Nusser and colleagues (37). We note that the reprogramming of E12.5 cTEC-progenitors to bipotency upon overnight culture indicates that these cells, although cTEC-fated, are not yet fully committed to the cTEC lineage and therefore we suggest they are regarded as cTEC-primed progenitors.

The mechanisms regulating the cTEC/mTEC cell fate decision and stabilization of cTEC and mTEC fates are currently unclear. Our findings extend the identification of Notch signalling as required for generation of the earliest mTEPC (33, 34) to implicate SOX9 as a potential regulator of early mTEC development. Further studies are required to address this possibility. While Notch signalling is essential for emergence of mTEC progenitors (33, 34), it does not explain the spatial organisation of the thymus into separate cortex and medulla regions, or their relative positioning. The interplay between RELB and Notch is unclear but may involve suppressing HES6 and thereby activating Notch in a similar way to neural stem cell maintenance by NF-κB (see Figure 1). Further work is required to elucidate these mechanisms, including how Notch signalling is limited in cTEC given their expression of Notch ligands such as DLL4 and Notch receptors such as Notch2. The possibility that cis-inhibition, through DLL4 expression, may play a role in cTEC fate restriction merits further investigation.

## Conclusion

The data presented herein support a revised model of the early events in thymic epithelial lineage development by providing phenotypic and functional evidence for the presence of cTEC- and mTEC-sublineage restricted progenitors from E11.5 and at least as early as E12.5, respectively. These findings, and the finding that cTEPC are reprogrammed to bipotency upon *in vitro* culture advance our understanding of early thymus development. They therefore have significance both for understanding of regulation of TEC progenitors later in ontogeny and for clinically-related thymus research, including research aimed at producing functional TEC from pluripotent stem cells (PSC) or by reprogramming. Our data suggest that establishment of fully defined medium able to support different TEC differentiation states, and of cTEPC and mTEPC-specific differentiation conditions, may expedite progress in this highly important research area.

## Disclosure of Potential Conflicts of Interest

The authors declare that the research was conducted in the absence of any commercial or financial relationships that could be construed as a potential conflict of interest.

## Author contributions

**AC, AMF, SP:** Conception and design, collection and assembly of data, data analysis and interpretation, manuscript writing. **DL, PR:** Conception and design, collection and assembly of data, manuscript writing. **VM, RM, JN, JS, JU:** Collection and assembly of data. **AIK, JM, SRT, JC:** data analysis and interpretation. **LB:** Provision of study materials. **CCB:** Conception and design, financial support, provision of study material, data analysis and interpretation, manuscript writing, final approval of manuscript.

## Data Sharing Statement

The datasets generated for this study can be found in GEO: GSE232765 and GSE229248. The codes for the analyses shown in Figures 1-3 and 7 and related supplementary data are available at: https://github.com/purewaltzan/ThymusDevelopment.

## Funding

The research leading to these results received funding from the School of Biological Sciences, University of Edinburgh (AC, DL), the UKRI-Biotechnology and Biological Sciences Research Council through an industrial CASE studentship (award number BB/MO16412/1; CCB, PR), the Darwin Trust (JM), the European Union Seventh Framework Programme (FP7/2007-2013) collaborative project ThymiStem under grant agreement number 602587 (CCB, AIK, SRT, DL), Leukaemia Research Fund Bennett Fellowship 0101 (CCB, AMF) and LRF Senior Lecturership 06008 (CCB, AMF), the UKRI-Medical Research Council through a DTP in Precision Medicine studentship (award number MR/N013166/1) (JS), and the Wellcome Trust collaborative award 211944/Z/18/Z (CCB, PR, SP, JC).

## Acknowledgements

We thank S. Monard, C. Cryer and F. Rossi (CRM, University of Edinburgh) for cell sorting, D. Kelly and M. Vermeren (CRM, University of Edinburgh) for imaging, and the BVS staff for animal care.

## Notes

### Competing Interest Statement

The authors have declared no competing interest.

