## Supplementary Information and data for "Thymic epithelial cell fate and potency in early organogenesis assessed by single cell transcriptional and functional analysis"

### Supplementary Figures

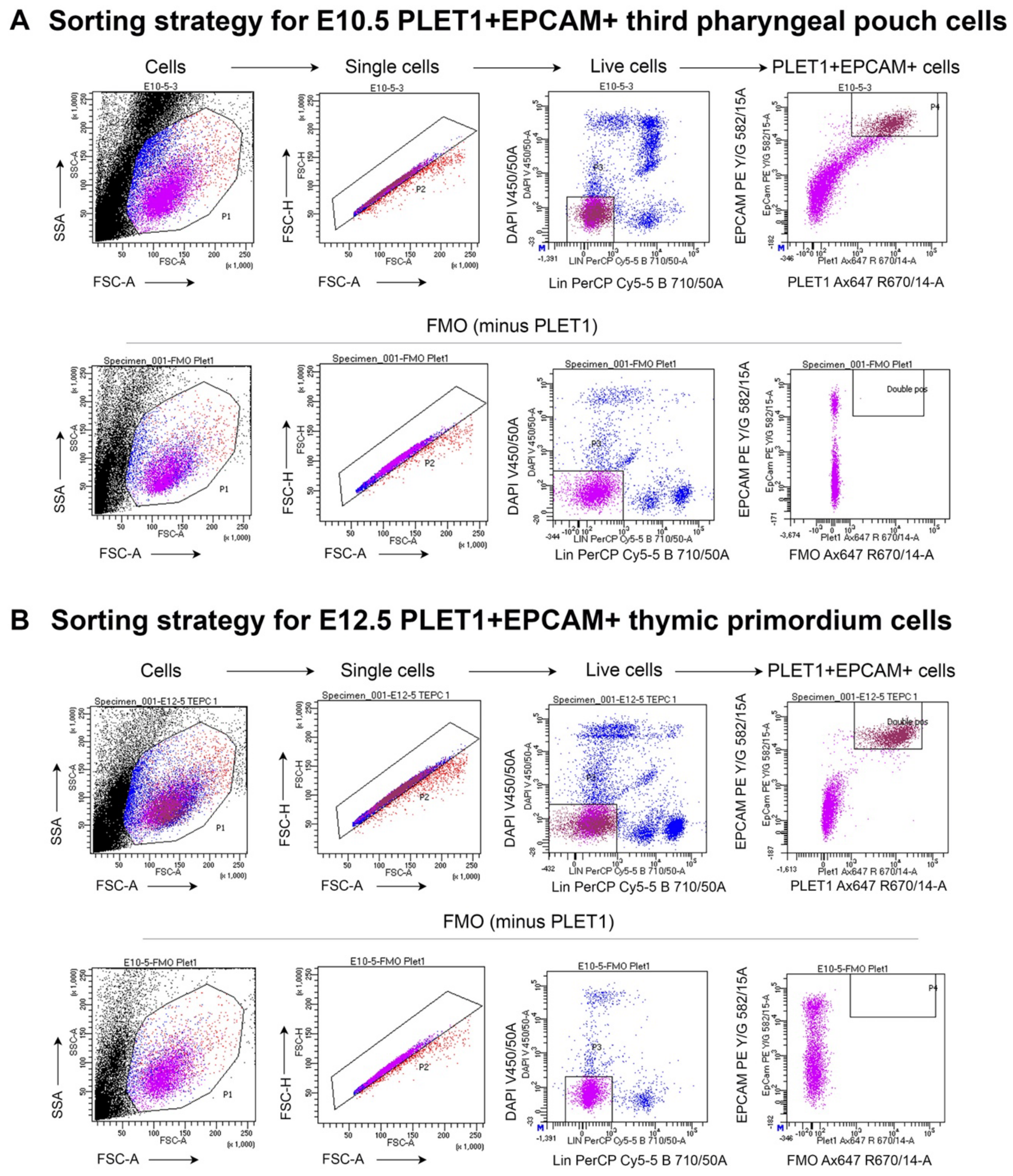


**Supplementary Figure 1. Fluorescence Activated Cell Sorting Strategy for E10.5 and E12.5 scRNAseq experiment.** Images shows representative plots from A) E10.5 third pharyngeal pouch cells and B) E12.5 thymic primordium cells. A) E10.5 third pharyngeal pouches plus some surrounding tissue and B) E12.5 thymic primordia were microdissected from mouse embryos and processed for fluorescence activated cell sorting as described in Materials and Methods. Sort gates were set using compensation control beads and FMOs. The sorting strategy was as shown. Lin = $\alpha$–CD45, $\alpha$–PDGFR$\alpha$, $\alpha$–PDGFR$\beta$.

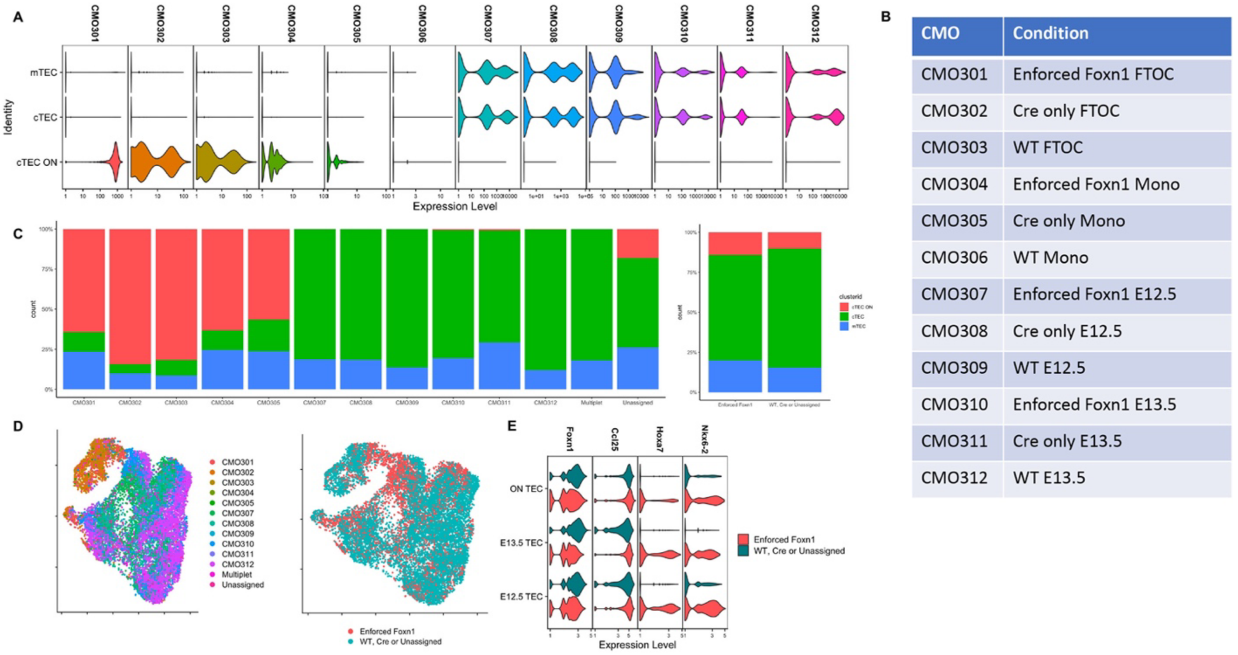


| CMO | Condition |
|---|---|
| CMO301 | Enforced Foxn1 FTOC |
| CMO302 | Cre only FTOC |
| CMO303 | WT FTOC |
| CMO304 | Enforced Foxn1 Mono |
| CMO305 | Cre only Mono |
| CMO306 | WT Mono |
| CMO307 | Enforced Foxn1 E12.5 |
| CMO308 | Cre only E12.5 |
| CMO309 | WT E12.5 |
| CMO310 | Enforced Foxn1 E13.5 |
| CMO311 | Cre only E13.5 |
| CMO312 | WT E13.5 |

**Supplementary Figure 2. Enforcing Foxn1 has only a modest effect on cell fate. (A)** Cells from different experimental conditions were tagged with different CMOs (see Materials and Methods). Some CMOs had very low counts. (**B**) Table showing CMO IDs for different experimental conditions. (**C,D**) Enforcing *Foxn1* only had a modest effect on cell fate, as seen from cluster distributions (**C**) or on UMAP plots (**D**). (**E**) Enforcing *Foxn1* increased expression of some genes, such as *Ccl25*, *Hoxa7* and *Nkx6-2*, but decreased expression of endogenous *Foxn1*.

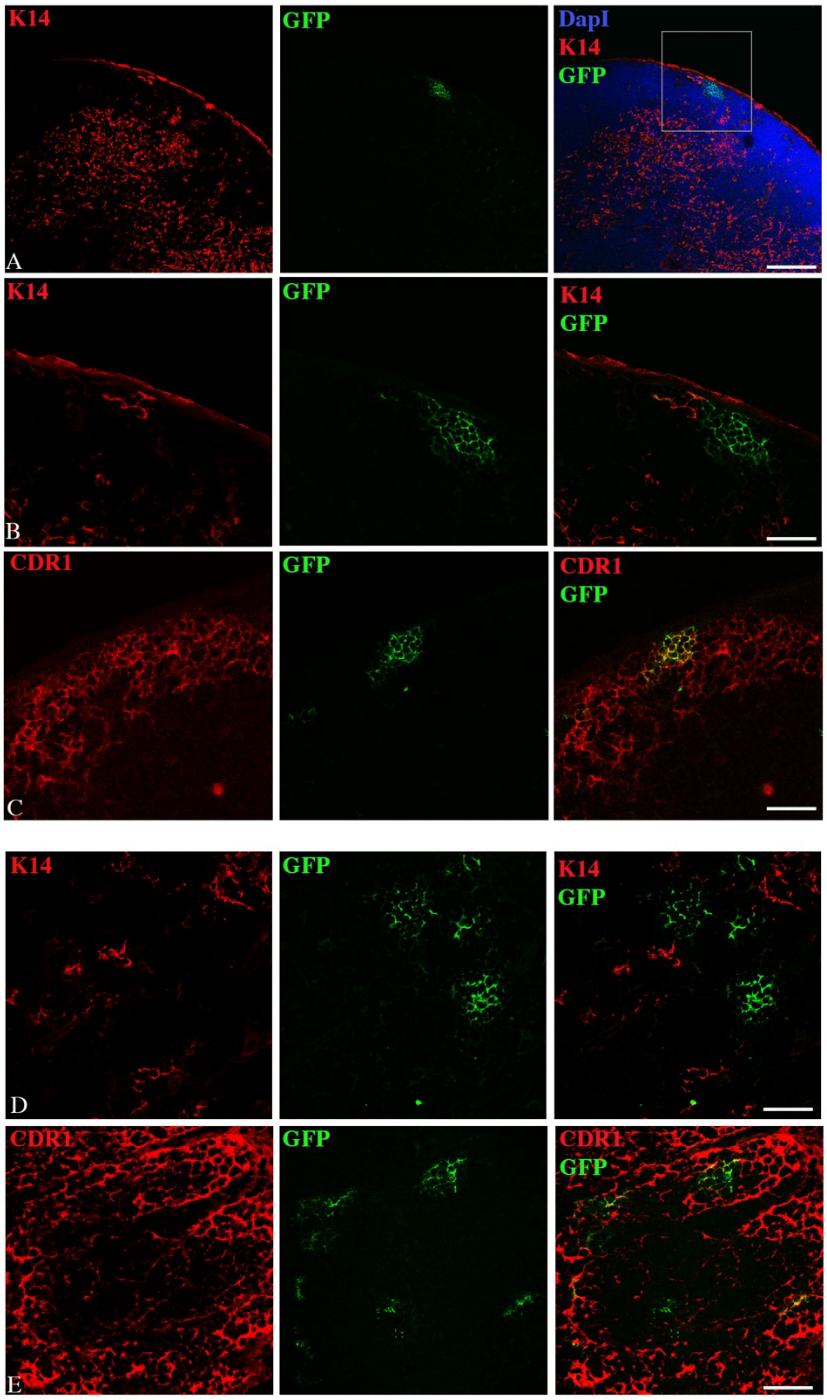


**Supplementary Figure 3. Sorting on EPCAM rather than PLET1 does not alter the outcome of the potency analysis.** Forty E12.5 mGFP$^+$EPCAM$^+$ cells were injected into an E12.5 thymic lobe, grafted under the kidney capsule and recovered after 2-3 weeks. Images show immunostaining of frozen sections with anti-cytokeratin 14 (K14) and CDRI. (**A-C**) show images of the same region of the recovered graft with a mGFP$^+$ foci within the cortical region expressing the mature cortical marker CDRI. (**D and E**) show other mGFP$^+$ foci within the same graft expressing the CDRI antigen. Scale bars, **A**=150μm, **B**=75μm, **C,D**=55μm

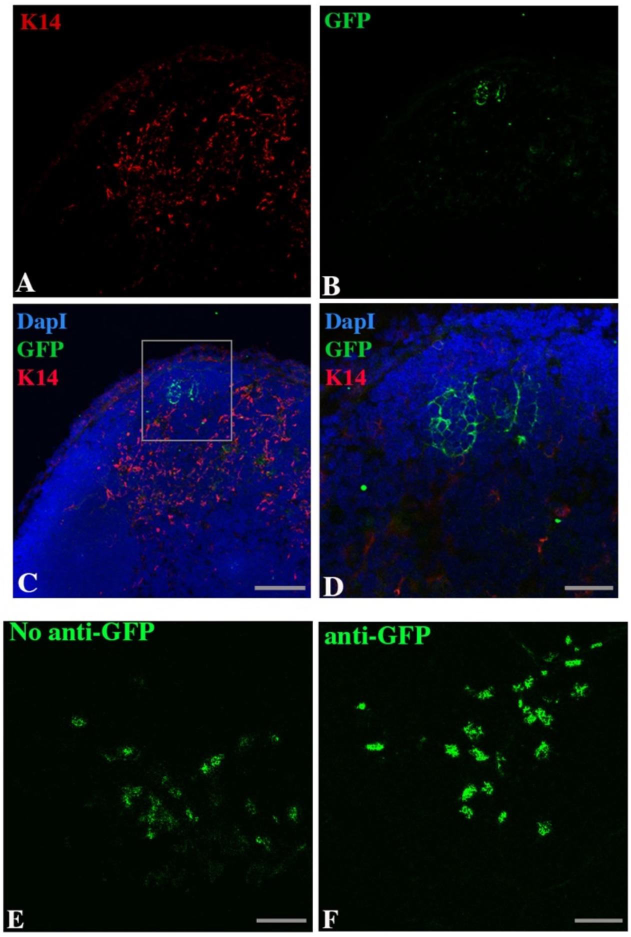


**Supplementary Figure 4. Altering the original protocol does not change the contribution or distribution of mGFP$^+$ progeny.** Images show immunostaining on frozen sections using $\alpha$-K14 and the nuclear stain DAPI. Changing the strain of mice from which the mGFP$^+$ cells were collected to CBAxC57BL/6 F1 (from C57BL/6), such that they were syngeneic with the host mice (CBAxC57BL/6 F1), made no difference to the results, most grafts contained clusters of mGFP$^+$ cells localised to only to cortical regions (**A,B**). (**E,F**) images show 488 fluorescence without (**E**) and with (**F**) $\alpha$-GFP antibody. Although using the $\alpha$-GFP antibody enhanced the GFP signal, the same number of clusters and distribution was observed. Scale bars, **A-C**, **E, F** = 150$\mu$m, **D** = 55$\mu$m.

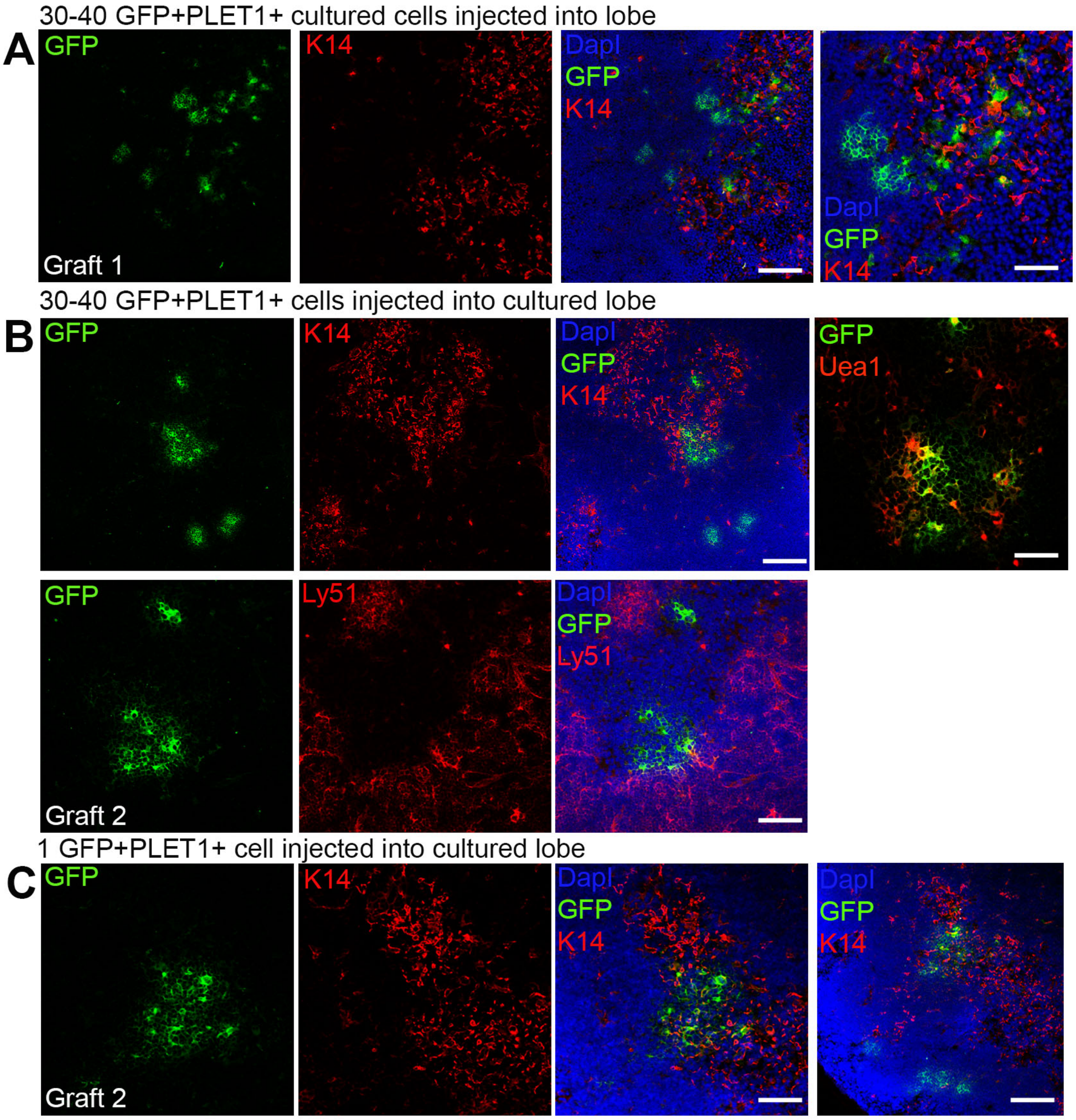


**Supplementary Figure 5. Culturing cells before injection or culturing lobes prior to injection results in increased medullary region contribution of mGFP$^{+}$ TEC**. Images show frozen sections of recovered grafts after either the injected E12.5 mGFP$^{+}$PLET1$^{+}$ TEC had been cultured overnight or the E12.5 lobes to be injected had been cultured overnight in DMEM, 10% FCS. (**A**) 30-40 cells cultured over night in DMEM, 10% FCS before injection showed a contribution to medulla in all recovered grafts. No grafts had contribution just to cortex. (**B, C**) Culturing lobes over night before injection of mGFP$^{+}$PLET1$^{+}$ TEPCs, resulted in medullary contributions in all grafts recovered. Graft name corresponds with graft identification in Table 3. Scale bars, A 75μm, and 55μm; B 300μm, 55μm and 75μm; C 75μm and 300μm.

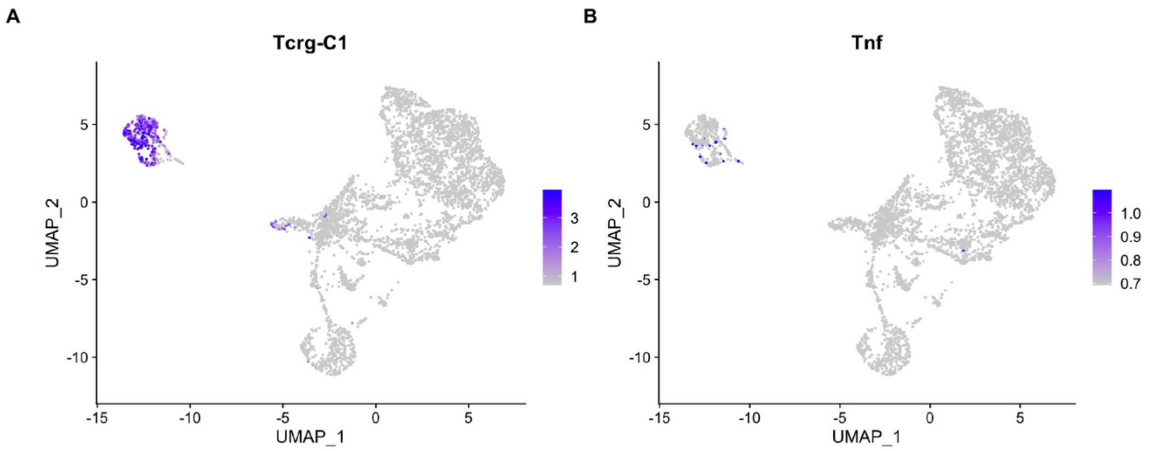


**Supplementary Figure 6. T-cells from the overnight culture condition express *Tnf*. (A,B)** UMAP plots of cells from the overnight culture condition, including TEC, parathyroid cells, fibroblasts and thymocytes. Thymocytes can be identified with the marker gene *Tcrg-C1* (**A**) and can also be seen to express some *Tnf* (**B**), which may be the source of the NF-kappaB signaling after overnight culture.

### Supplementary Tables

| *Transgene* | *Orientation* | *Sequence* |
|---|---|---|
| Foxn1$^{Cre}$ | Forward | 5’ GAC CAG GTT CGT TCA CTC ATG G |
| Foxn1$^{Cre}$ | Reverse | 5’ CCT TAG CGC CGT AAATCA ATC G |
| Sox9CreER$^{T2}$ | Forward | 5’ CGG TTT CGT TCT CTG TTT TCC |
| Sox9CreER$^{T2}$ | Reverse | 5’ AGG CAA ATT TTG GTG TAC GG |
| mGFP | Forward | 5’ ACA TGG TCC TGC TGG AGT TC |
| mGFP | Reverse | 5’ TCA GGT TCA GGG GGA GGT |
| Rosa26-CreERT2 | Forward | 5’ GCA TAA CCA GTG AAA CAG CAT TGC TG |
| Rosa26-CreERT2 | Reverse | 5’ GGA CAT CAG GGA TCG CCA GGC G |
| iFoxn1 | Forward | 5’ GGG AGC AGC TGA AGG ATG AC |
| iFoxn1 | Reverse | 5’ CGC TTG AGG AGA GCC ATT TG |

**Supplementary Table 1.** Primers used to genotype transgenic lines. Note, Gt(ROSA)26Sor$^{tm14(CAG\text{-}tdTomato)Hze}$ (Ai14) were maintained as homozygotes.

| Antibody | Clone | Supplier | Staining |
|---|---|---|---|
| α-Beta5t | Polyclonal | MBL International | IHC |
| CD205 | NLDC-145 | AbD Serotec | IHC |
| CDR1 | CDR1 (rat IgG2a) | B. Kyewski | IHC |
| Ly-51 | 6C3 (rat IgG2a) | Biolegend | IHC |
| α-Cytokeratin 14 (LL002) | Polyclonal | B. Lane, University of Dundee | IHC |
| α-Cytokeratin 14 (AF 64) | Polyclonal | Covance | IHC |
| α-Cytokeratin 8 | Troma1 | DSHB | IHC |
| α-Pan-Cytokeratin | Polyclonal | DAKO | IHC |
| UEA-1 biotin | Lectin | Vector Labs | IHC, FC |
| α-SOX9 | Rabbit anti-mouse | Cell Signaling Technology | IHC |
| α-GFP-488 | | Molecular Probes | |
| α-EPCAM-APC | G8.8 (rat IgG2a) | BioLegend | FC |
| α-EPCAM | G8.8 (rat IgG2a) | DSHB | FC |
| α-EPCAM-FITC | 9C4 | Biolegend | FC |
| α-MHC Class II PE-Cy7 | M5/114.15.2 | BioLegend | FC |
| α-PLET1 | 1D4 (rat IgG) | In house | FC |
| α-PLET1 | MTS20 (rat IgM) | R. Boyd, Monash University | FC |
| α-CD45 | | BD Biosciences | FC |
| DAPI | | Invitrogen | Viability dye |
| DAPI | | Life Technologies | Viability dye |
| Goat anti-rabbit Alexa Fluor$^{TM}$ 647 | Polyclonal | Invitrogen A48265 | Secondary antibody IHC |
| Goat anti-rabbit Alexa Fluor$^{TM}$ 568 | Polyclonal | Invitrogen A11077 | Secondary antibody IHC |
| Streptavidin-FITC | | Vector Labs B1065 | Secondary reagent IHC |

**Supplementary Table 2.** Antibodies used Panel for flow cytometry and immunohistochemistry. Table shows the name, associated fluorophore where appropriate, supplier and clone number. DAPI, (4',6-Diamidino-2-Phenylindole, Dihydrochloride); IHC, immunohistochemistry; FC, flow cytometry.

| *Antibody* | *Barcoded Reagent* | *Dilution* | *Supplier* |
|---|---|---|---|
| UEA1-Biotin | TotalSeqTM-B0952 Streptavidin-PE | 1/1,500 | Biolegend |
| MHCII ($\alpha$-mouse I-A/I-E) | TotalSeq$^{TM}$-B0117 | 1/10,000 | Biolegend |
| $\alpha$-CD40 | TotalSeq$^{TM}$-B0903 | 1/3,200 | Biolegend |
| $\alpha$-CD80 | TotalSeq$^{TM}$-B0849 | 1/3,200 | Biolegend |
| $\alpha$-EPCAM | Beads | 1/10 | Miltenyi |

**Supplementary Table 3.** Cell staining reagents used in 10x sequencing experiment.

**Supplementary Tables 4 and 5** are provided as separate files.

**Supplementary Table 4.** GO analysis comparing the clusters "ON cTEC" and "cTEC" from E12.5, E13.5 and ON TEC. GO terms that are enriched in "ON cTEC" are shown in columns B-L, while GO terms that are depleted are shown in columns M-W.

**Supplementary Table 5.** Differential gene expression analysis comparing the clusters "ON cTEC" and "cTEC" from E12.5, E13.5 and ON TEC. Columns meanings are as follows: p_val: p-value for differential expression based on wilcox test; avg_logFC: log fold-chage of the average expression between the two groups. Positive values indicate that the gene is more highly expressed in the first group; pct.1: The percentage of cells where the gene is detected in the first group; pct.2: The percentage of cells where the gene is detected in the second group; p_val_adj: Adjusted p-value, based on bonferroni correction using all genes in the dataset.
